# Cell class-specific electric field entrainment of neural activity

**DOI:** 10.1101/2023.02.14.528526

**Authors:** Soo Yeun Lee, Konstantinos Kozalakis, Fahimeh Baftizadeh, Luke Campagnola, Tim Jarsky, Christof Koch, Costas A. Anastassiou

## Abstract

Electric fields affect the activity of neurons and brain circuits, yet how this interaction happens at the cellular level remains enigmatic. Lack of understanding on how to stimulate the human brain to promote or suppress specific activity patterns significantly limits basic research and clinical applications. Here we study how electric fields impact the subthreshold and spiking properties of major cortical neuronal classes. We find that cortical neurons in rodent neocortex and hippocampus as well as human cortex exhibit strong and cell class-dependent entrainment that depends on the stimulation frequency. Excitatory pyramidal neurons with their typically slower spike rate entrain to slow and fast electric fields, while inhibitory classes like Pvalb and SST with their fast spiking predominantly phase lock to fast fields. We show this spike-field entrainment is the result of two effects: non-specific membrane polarization occurring across classes and class-specific excitability properties. Importantly, these properties of spike-field and class-specific entrainment are present in cells across cortical areas and species (mouse and human). These findings open the door to the design of selective and class-specific neuromodulation technologies.

## Introduction

Delivery of exogenous electric fields of various shapes and forms through stimulation devices to alter brain activity has a rich history in neuroscience. Since the early work by Fritsch and Hitzig mapping brain region during surgery ^1,2^, extracellular electric stimulation (ES) has been used to study functional connectivity, excitability and response properties of the brain. In therapeutic interventions, ES alleviates symptoms of neurological disorders such as Parkinson’s disease^3–5^, dystonia^6^, depression^7–9^, epilepsy ^10,11^ and others (e.g. see review ^12^). Despite these advances, there is still a lack of understanding at the basic cellular level of what happens when brain matter is electrically stimulated to promote certain activity patterns or suppress others ^13–17^. This has limited the application of ES as a basic science tool and, importantly, as a therapeutic intervention for brain disorders.

Brain circuits consist of multiple distinct and diverse cell classes yet how ES affects these classes individually at a cellular level remains unknown. Most studies of ES focus on the excitatory glutamatergic neurons in various brain regions (neocortex ^18,19,17^; hippocampus ^20–23^; but also ^24–26^). While inhibitory GABAergic cells are less in absolute numbers, they play critical roles in structuring and regulating brain circuit activity ^27–31^. Given such diversity, whether and how ES impacts excitatory and inhibitory classes remains unclear, as well as the extent to which cell classes may respond distinctly to ES. For example, it is unclear whether inhibitory classes entrain to the ES like excitatory neurons, or whether they remain unperturbed due to their morphological compactness ^16,32^. As a result, ES protocols applied in the brain of both animal models and humans do so without consideration for the remarkable cellular diversity comprising brain circuits.

Here we study the direct impact of extracellular ES in two cortical brain regions, visual cortex and CA1 hippocampus, using simultaneous intracellular (whole-cell patch-clamp) and extracellular recordings from identified single neurons of major excitatory and inhibitory cell classes. We observe strong, differential and cell class-specific spike-field entrainment with excitatory neurons entraining already at slow ES frequencies (8 Hz) while inhibitory neurons only entraining for fast ES frequencies (30-140 Hz). We also show that the class-dependence of spike entrainment is not the result of a differential subthreshold (i.e., non-spiking) mechanism as cortical neurons across classes, brain regions and, in fact, species exhibit ubiquitous, robust, and frequency-independent membrane coupling to ES oscillations. The fact that we see similar entrainment properties in human excitatory cortical neurons suggests widely present biophysical mechanisms in which the unique ion-channel composition of each cell class dictates the class-specific entrainment to ES. The latter opens the door to the design of new ES waveforms and delivery systems offering improved selectivity and neural control.

## Results

To simultaneously stimulate and record intracellular and extracellular signals close to identified neurons, we use an *in vitro* experimental setup to examine the cellular and cell type-specific effects of the electric stimulation (**ES**)-generated field, at multiple locations around the cell body. To position multiple stimulation and recording sites in specific locations near a cell body we used a customized 8-pipette multi-patch brain slice electrophysiology system. It enables simultaneous extracellular stimulation, intracellular current injection, whole-cell patch-clamp recordings, and extracellular recordings from multiple sites within 50-120 µm of the soma (Figure 1a). ES effects are assessed in brain slices at the single compartment (soma) and membrane level, *i.e.*, on the extracellular (V_e_), intracellular somatic (V_i_), and the resulting membrane voltage (V_m_, calculated as V_m_=V_i_-V_e_) in the vicinity of the whole-cell patch-clamped soma. To study the impact of ES on V_e_, V_i_ and V_m_ of single neurons, we first estimated the electric field and resulting V_e_-deflection in the vicinity of the cell body from sinusoidal ES delivery through an extracellular pipette as a function of distance (Figure 1a-c). The highest ES amplitude (200 nA) induced a V_e_-amplitude of 1.1±0.5 mV (mean ± std) and a field amplitude of 22.6±8.9 mV/mm (mean ± std) close to the cell body (measured 15 μm from the soma). The V_e_-distance relationship was accurately captured by the point-source approximation for a purely resistive and homogeneous medium ^19,33^. We conclude that the ES amplitudes imposed in our study are relatively weak and within the physiological range measured in rodents and humans *in vivo* (*e.g.* ^16,34^).

**Figure 1:**
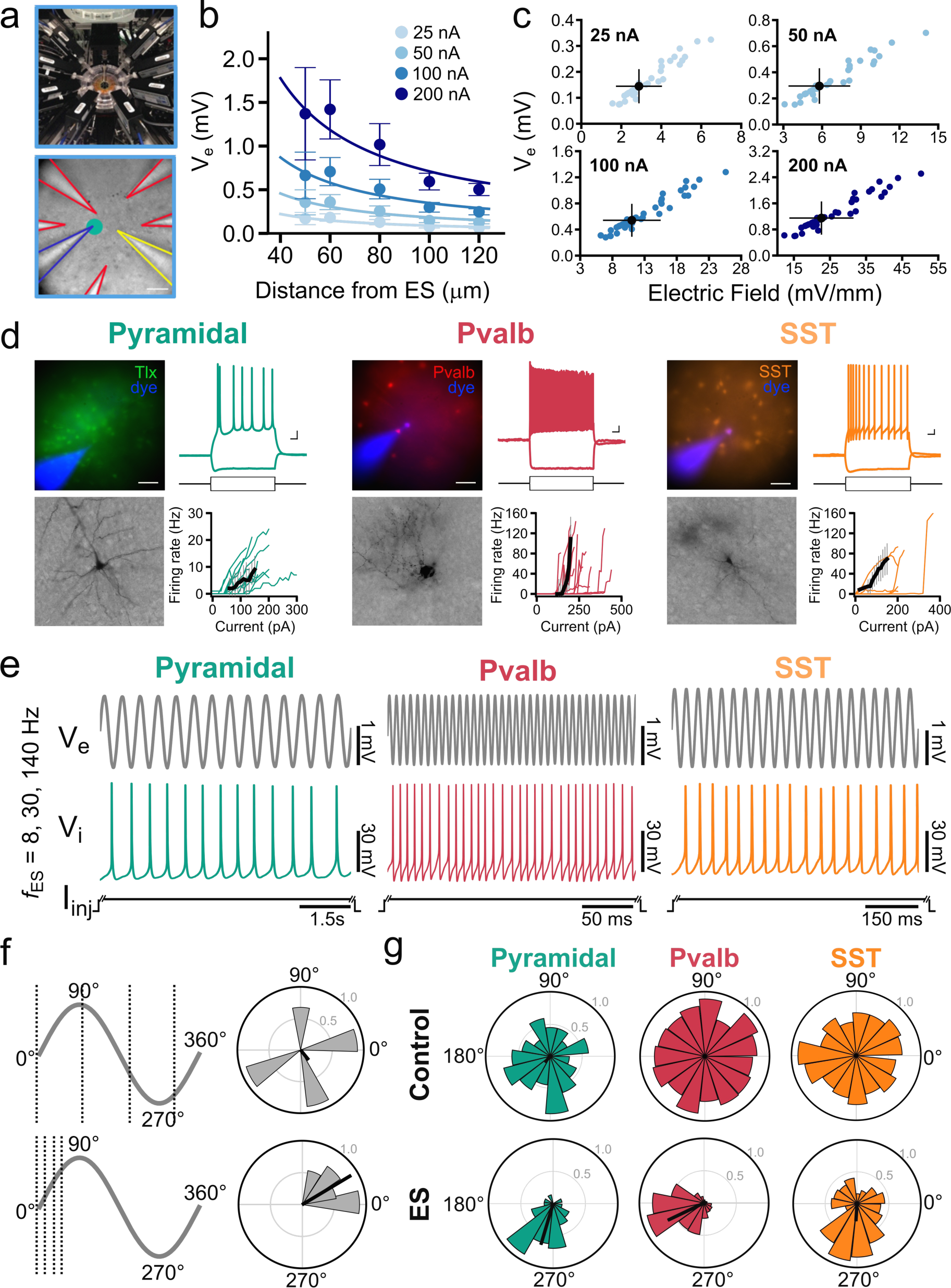
Cellular and cell type-specific characterization of ES effects. **a**, Top: Simultaneous ES, V_i_ and V_e_ recordings from eight electrodes near the soma of a whole-cell patched neuron. Bottom: Experiment in mouse cortical slice (yellow: extracellular ES, green: patched cell soma of layer 5 pyramidal neuron; blue: intracellular electrode recording V_i_; red: extracellular electrodes recording V_e_ at multiple locations within 150 um from soma). **b**, Left: V_e_ amplitude as function of distance between ES and recording electrodes (ES: amplitude is 25-200 pA; ES frequency: 8 Hz; circles: V_e_ mean amplitude; error bars: V_e_ amplitude st.d.) Trendlines are least-squares fit of the point source approximation. **c**, V_e_ and electric field amplitude elicited by the ES at the extracellular recording electrode closest to the whole-cell patched soma (approx. 15 µm). Blue: V_e_ amplitude for each experiment (n=59); Black: mean and st.d. across experiments. **d**, Sample fluorescent images, cellular morphology, electrophysiology responses, and response curves (spike frequency vs. current input, f-I) from identified neocortical cells (colored lines: individual neurons; black: median). Three cell types: (left) excitatory pyramidal, (middle) inhibitory Pvalb, and (right) inhibitory SST. **e**, Experiments with simultaneous ES (delivered at 8, 30, and 140 Hz for all cells) and intracellular DC current injection I_inj_ to elicit spiking. Sample electrophysiology traces from three cells. top, V_e_ close to the soma; bottom: V_i_ from inside the soma of a spiking neuron responding to I_inj_ (5 s of I_inj_ shown here). Left to right: traces from a pyramidal, a Pvalb and an SST neuron. **f**, Introduction of spike time analysis and quantification of spike-field entrainment. Spikes are ascribed a phase by mapping the spike time to the V_e_ phase from the electrode closest to the spiking cell. Spike phase distributions shown in polar plots. **g**, Top: Polar plots of the spike-phase distribution for a pyramidal cell (green), Pvalb cell (red), and SST cell (orange) in the absence (Control) and presence of ES (200 pA; 8 Hz, 140 Hz, 30 Hz ES for pyramidal, Pvalb, and SST cells, respectively). The population vector length (black line) reflects the degree of entrainment. The presence of ES skews the spike-phase distribution and entrains the neurons. Bottom: Spike phase distributions under ES for a pyramidal (green; ES: 200 pA, 8 Hz), a Pvalb (red; ES: 200 pA, 140 Hz) and an SST neuron (yellow, ES: 200 pA, 30 Hz). # of spikes, Pyr: Control: 52, ES: 56; Pvalb: Control: 702, ES: 864; SST: Control: 216, ES: 218.

We used acute slices of mouse primary visual cortex (V1) to focus on excitatory pyramidal cells and the two largest GABAergic cell classes in cortical layer 5 (L5), the parvalbumin (Pvalb), and somatostatin (SST)-expressing cells comprising approximately 54% and 25% of the Layer 5 (L5) GABAergic cell population, respectively^35^ (Figure 1d). We used several modalities (fluorescent markers in transgenic animals, electrophysiological features, and cellular morphology) to identify individual neurons from the three cell classes (Figure 1d). Specifically, excitatory pyramidal neurons were initially identified in Cre-or Flp-dependent transgenic mice (driven by L5-specific Tlx3 or Sim1 promoters) with fluorescent reporters for L5 excitatory neurons or by their pyramidal morphology, and then confirmed by their characteristic regular-spiking firing pattern in response to intracellularly injected current steps and slower F-I curve in patch-clamp recordings (e.g. ^36^). Inhibitory Pvalb and SST neurons were identified using Cre-or Flp-dependent transgenic mice with fluorescent reporters and confirmed by their fast-spiking firing pattern and steep F-I curve (Figure 1d) ^36^. The recorded cells were filled with biocytin, for *post-hoc* processing to reveal their cellular morphology and confirm the classification.

In principle, even weak ES can impact the cellular membrane (e.g. ^19^) but whether and how it does so for different cell classes remains speculative. To address this question, we stimulated (via intracellular DC current injection) and recorded from pyramidal, Pvalb, and SST neurons while delivering ES through an extracellular stimulating electrode placed 50 µm away from the cell body. Intracellular current injections of up to two-fold rheobase were used to elicit spiking during control and ES (Table S1; 154.5 ± 64.9 pA for pyramidal neurons; 259.5 ± 99.8 pA for Pvalb; 191.5 ± 102.8 pA for SST). The ES was delivered as a sinusoidal current at amplitudes from 25 to 200 nA. Such weak ES (Figure 1a-c) never elicited action potentials in the absence of other intracellular inputs ^18,19^. Furthermore, imposing the maximum ES amplitude caused no change in spike frequency in any of our experiments (Figure S1). The ES frequency was varied from 8 to 140 Hz, covering a range used in ES applications (Figure 1e). We then mapped the spikes during sinusoidal ES to their respective spike phase along the sinusoidal ES (reference signal: V_e_; Figure 1f). In a second step, spike-field entrainment of the different cell classes was assessed via population vector analysis (Figure 1g) (see also Methods). Increased spike-field entrainment due to the ES-induced field results in an inhomogeneous spike phase distribution with a distinct, preferred spike phase and, therefore, a larger population vector. A weak ES effect on neural spike timing or decreased entrainment, in turn, is marked by a homogeneous spike phase distribution and a small population vector length (e.g., control experiments where no ES is delivered; Figure 1g). We found a diversity in the entrainment profile of cortical cell classes that depends on ES parameters as well as cell class properties and activity (Figure 1g).

### Rich, diverse and cell class-specific spike-field entrainment to ES

To understand the origin of differential electric field entrainment, we first looked at the influence of ES on the spiking cellular membrane. Hitherto, the assumption has been that ES has the same effect on the neural membrane across the various classes as long as experimental parameters (e.g., distance of the ES electrode, the induced field, etc.) are controlled for. To test this hypothesis, we conducted a set of experiments where we analyzed the spiking activity of cells in the presence vs. absence of ES. To isolate the pure ES effect from effects mediated from other neurons, experiments were conducted under synaptic blockers (see Methods). To elicit spiking, we injected intracellular DC current into the cell body of neurons (intracellular stimulation duration: 9 s). This was repeated under two conditions: in the absence of any ES (“control”) and, in a next step, the same intracellular current was injected while concurrently applying ES at various frequencies (*f_stim_* = 8, 30, 140 Hz) and amplitudes (25 to 200 nA) (Figure 2a).

**Figure 2.**
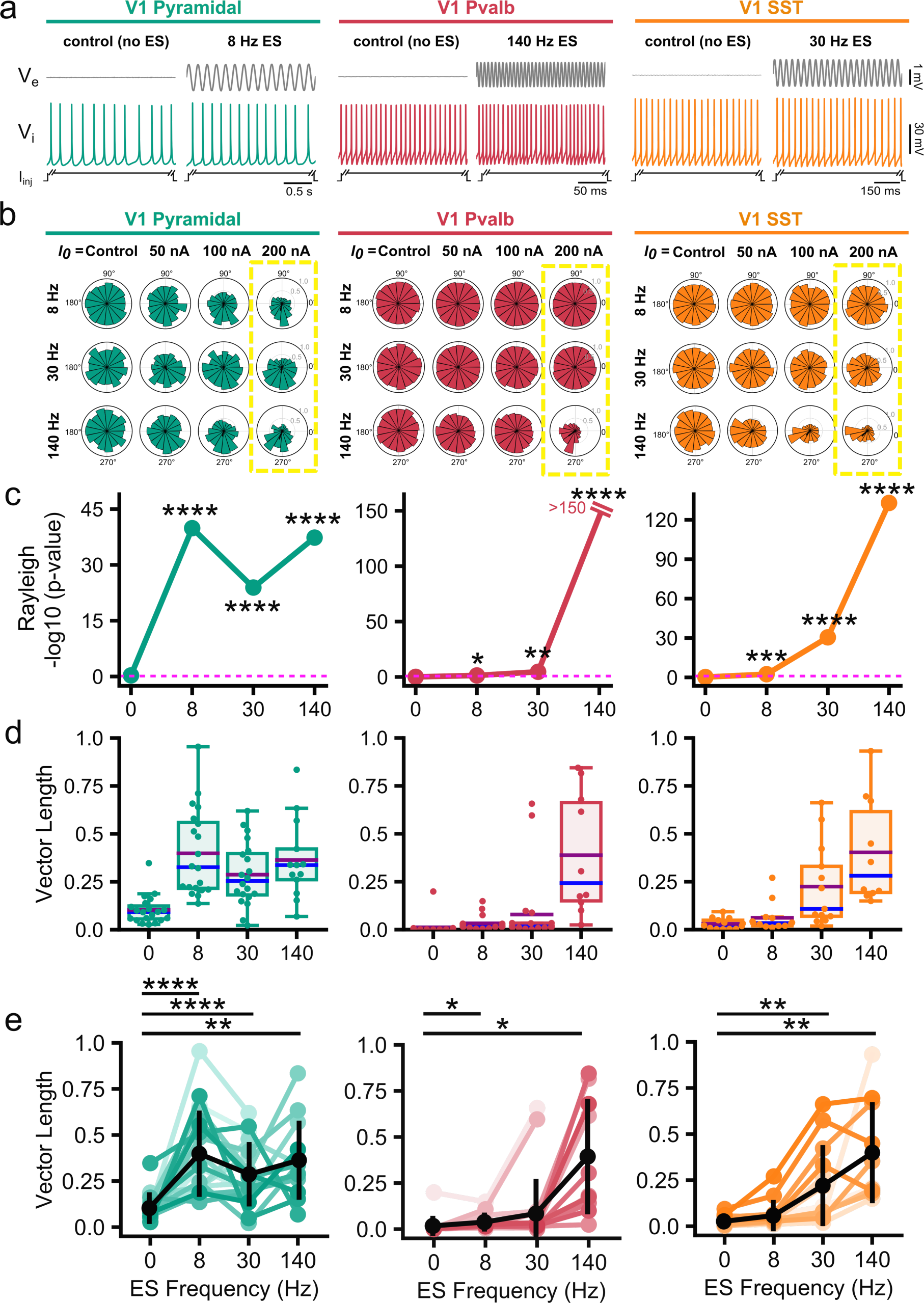
Cell class-specific entrainment of spiking to ES. **a,** Simultaneous spiking induced by a suprathreshold DC stimulus I_inj_ co-occurs with sinusoidal ES. Despite its relatively small amplitude (see Figure 1c and Figure 2), the subthreshold effect entrains neuronal spike timing (left to right: traces from a pyramidal, a Pvalb and an SST neuron) without affecting the spike rate (see Figure S1). **b**, Spike-phase distribution for pyramidal (green), Pvalb (red), and SST (orange) cell classes for varying ES parameters. (Rows) ES frequency (top to bottom): 8, 30, to 140 Hz. (Columns) ES amplitude (left to right): 0 (Control), 50, 100 and 200 nA. “Control” indicates no ES is applied (ES amplitude: 0 nA). Control experiments show symmetric spike-phase distribution, i.e., no inherent preferred spike-phase in the absence of ES, as assessed by Rayleigh’s test (p>0.05, see Table S2). Increasing (but subthreshold) ES amplitude generally increases spike-phase entrainment. Different ES frequencies have a distinct, cell class-specific effect on spike phase entrainment. Summary statistics for the highest amplitude (in yellow box) plotted in **c**. **c-e,** Spike entrainment to ES assessed via the Rayleigh’s test (**c**) and the population vector length (**d-e**) against the ES frequencies (ES amplitude: 200 nA). **c,** p-values calculated for pyramidal, Pvalb, SST spikes (left to right) via Rayleigh’s test as function of ES frequency show statistically significant spike phase entrainment (*p ˂ 0.05, **:p ˂ 0.01, ***p ˂ 0.001, ****p ˂ 0.0001, see Table S2). Dashed pink line: p-value at 0.05. Pyramidal spiking entrains to slow (8 Hz), medium (30 Hz), and fast (140 Hz) ES frequencies while Pvalb and Sst are most strongly entrained by high ES frequencies (140 Hz). **d**, The population vector length plotted for each cell (gray circles) within each class (left to right: Pyramidal, Pvalb, SST). Purple and blue lines: mean and median values, respectively; error bars: st.d. **e**, The population vector length for each cell (from **d**) across each ES frequency is compared against control conditions (no ES) to assess degree of entrainment (paired t-test, false discovery rate (FDR)-corrected for multiple comparisons: *p<0.05, **p<0.01, ****p<0.0001) Pyramidal: N=21 cells for 8, 30 Hz, N=13 for 140 Hz; Pvalb: N=22 cells for 8, 30 Hz, N=12 for 140 Hz; Sst: N=13 for 8, 30 Hz, N=10 for 140 Hz.

To what extent does ES impact spiking? Notably, the presence or absence of weak ES did not alter the spike frequency elicited by the intracellular DC injection for all cell classes tested (Figure S1). We next tested whether ES affected the temporal patterning of spikes: does ES give rise to increased or decreased spike entrainment? Because the imposed ES and the recorded V_e_ in the proximity of the recorded cells is sinusoidal, spike times can be mapped to spike phases (Figure 1e-f; reference signal: V_e_ from the electrode closest to the recorded neuron; spike phase: 0° – 360°). To test whether ES altered the temporal patterning of spike times, we looked for an inhomogeneity in the spike phase distribution as function of cell class and ES parameters. To control for spurious correlations, we devised the following control experiment: adopt the V_e_-signal from the subsequent experiment where ES is present (i.e., ES amplitude is non-zero), map spike times to spike phases and run the same statistical tests for this “virtual” ES experiment. Under such conditions, a time-locked but ES-independent mechanism at any of the tested ES frequencies will result in an inhomogeneous non-uniform spike-phase distribution under control conditions. To assess whether the spike-phase distribution is non-uniform, we used circular statistics^19,21^. Specifically, the population vector length (range 0-1) and the Rayleigh test (null hypothesis: spike-phases are uniformly distributed along the unit circle and exhibit no preferred phase), serve as a metric of spike phase coordination. The lack of spike-phase coordination results in a uniform spike-phase distribution, a small vector length and a statistically insignificant *p*-value of the Rayleigh test. In contrast, the presence of spike-phase coordination due to ES gives rise to a non-uniform spike-phase distribution (with spike-phases mapped based on the local V_e_ signal dominated by the ES), a large vector length, and a statistically significant *p*-value of the Rayleigh test.

We first looked at pyramidal neurons, where it is known that longer current step injections lead to temporally irregular spiking ^19,37^. We saw that, indeed, for control experiments, homogeneous spike-phase distributions appear with very small vector lengths, i.e., no preferences in terms of spike-phase in the absence of ES (Figure 2b-c, Table S2). When we increased the ES amplitude, spike-phase entrainment appears, as seen in the decreasing *p*-value (calculated by the Rayleigh test) and increasing vector length (Figure 2c-e, ES amplitude: 200 nA; Figure S2, ES amplitude: 100 nA, Table S2). Importantly, while spike phase distribution changed with ES, the average spike frequency remained unaltered (Figure S1). We note that even at the lowest ES amplitude of 25 nA, resulting in a field strength of approx. 1-2 mV/mm and a V_e_-amplitude of 0.1 mV, strong spike-field coordination occurs across frequencies for excitatory and inhibitory classes (Figure S3). It follows that weak, subthreshold ES results in strong pyramidal spike-phase coordination despite leaving the spike rate unaffected. Furthermore, spike-field entrainment of pyramidal neurons is ubiquitous and reaches very high ES frequencies. Specifically, while Pyr exhibit a preferred spike-field entrainment at 8 Hz ES, the same pyramidal neurons also exhibit strong spike-field coordination at 140 Hz (Figure 2b-c). We conclude that weak, subthreshold ES results in strong pyramidal spike-phase coordination and that the entrainment remains strong even for fast ES frequencies above 100 Hz.

We next looked at the spike-field entrainment of major cortical inhibitory cell classes. We ran similar suprathreshold experiments (control vs. ES) with Pvalb and SST interneurons and performed identical analyses calculating the same entrainment metrics. In sharp contrast to pyramidal neurons, we found that both Pvalb and SST entrain their spiking predominantly to high ES frequencies: SST entrain for >30 Hz whereas Pvalb exhibit strong spike-field coordination for 140 Hz and much weaker coordination for slower ES of the same amplitude (Figure 2c-e, Table S2). Once more, the spike-field entrainment observed in Pvalb and SST is solely attributed to the impact of ES: control experiments where ES is absent (zero amplitude) show no coordination between spiking and any effect in a similar time scale to ES for any cell class. It is only when weak ES is imposed at the relevant frequency for each inhibitory class that Pvalb and SST spiking is entrained (e.g., compare Pvalb spike-phase distributions for 8 and 30 Hz; Figure 2b; Figure S2, S4).We conclude that while Pvalb and SST inhibitory neurons exhibit strong spike-field entrainment to ES, this entrainment depends on the ES parameters, with cells exhibiting a preference to fast ES (>30 Hz for SST and 140 Hz for Pvalb).

Surprised by the richness of the spike-field entrainment characteristics, we sought to reproduce our basic findings in another brain region. We repeated our experimental protocols in the hippocampus, recording from pyramidal neurons and Pvalb located in the CA1. We used the same ES protocols as in V1, which produced similar field strengths (Figure S5). As for V1 neurons, we found that in CA1 neurons weak ES causes strong pyramidal spike-phase coordination, resulting in increasing spike-field entrainment that remains strong even for fast ES frequencies (> 100 Hz; Figure S5-S6). Hippocampal inhibitory Pvalb on the other hand, like their V1 counterparts, exhibit a much more nonlinear spike-field entrainment profile with a clear preference to fast ES (Figure S5f-i, S6, Table S3). We conclude that the rich and diverse spike-field entrainment characteristics to ES shown in V1 are also observed in hippocampal CA1, indicating that our findings are robust and due to biophysical mechanisms shared across cortical areas.

### Ubiquitous and robust subthreshold entrainment across cell classes

Given the strong and class-specific entrainment of spiking to ES, we asked how this effect is mediated. Specifically, we asked whether ES also impacts the non-spiking (subthreshold) membrane in a class-specific manner that results in the observed spike entrainment. We analyzed pyramidal, Pvalb, and SST neurons by either holding them at their resting potential (Table S1) or at various subthreshold polarization levels and delivered sinusoidal ES of varying amplitude (25-200 nA) and frequency (1-100 Hz) at 50 µm from cell soma. To assess the effects of ES on the subthreshold neurons, we analyzed the resulting amplitude and phase deflections of V_e_, V_i_, and V_m_ (Figure 3).

**Figure 3.**
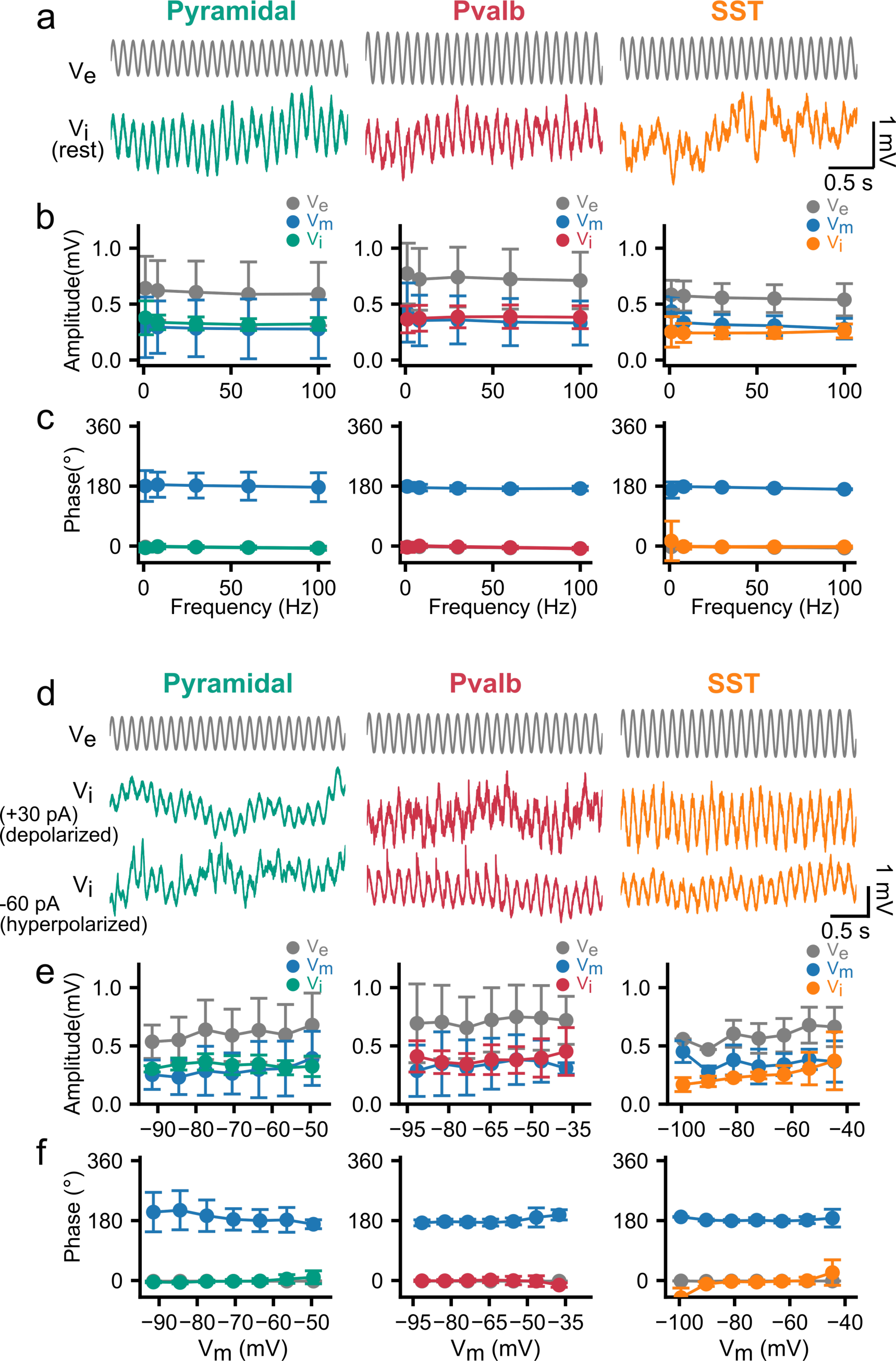
Non-specific, ES frequency-independent subthreshold entrainment of excitatory and inhibitory cell classes to ES. **a**, Entrainment of V_i_ to ES, for pyramidal (green), Pvalb (red), and SST (orange) cells. Gray traces: V_e_ measured at the closest extracellular location (15 µm) from the cell soma. The subthreshold sinusoidal ES is delivered 50 µm from cell soma (frequency: 8 Hz, amplitude: 100 nA). **b, c** ES effect on neurons at resting potential (ES amplitude: 100 nA, frequency: 1 to 100 Hz). Amplitude (**b**) and phase (**c**) of the ES where V_e_ (gray), V_i_ (green, red, or orange), and V_m_ (blue) for each cortical cell type (circles: mean; error bars: std; left to right: pyramidal, Pvalb, SST). The three cell classes exhibit ES frequency independence with induced V_i_, V_e_ and V_m_ amplitude and phase remaining constant for ES frequencies ranging 1-100 Hz (one-way ANOVA, p>0.05 for Pyr, Pvalb, and SST). **d**, ES effect on hyper- and depolarized neurons held at a range of membrane potentials via injection of depolarizing or hyperpolarizing current I_inj_ (from −90 to 90 pA). The sinusoidal ES is simultaneously applied with I_inj_ (ES amplitude: 100 nA, frequency: 8 Hz). **e, f** Amplitude (**e**) and phase (**f**) of the ES-induced V_e_ (gray), V_i_ (green, red, or orange), and V_m_ (blue) for each cortical class for hyper- and depolarizing I_inj_ (circles: mean; error bars: std; left to right: pyramidal, Pvalb; right: SST). The induced ES effect on V_i_, V_e_ and V_m_ amplitude and phase are broadly independent of membrane polarization and remain constant across a range of 40 mV for the three classes. N=24 cells (Pyramidal), N=22 cells (Pvalb), N=6 cells (SST).

In the presence of synaptic blockers (see Methods) and in the absence of intracellular current injection, the V_e_, V_i_, and V_m_ response (amplitude and phase) to ES measures the pure subthreshold ES effect on the cellular membrane (Figure 3a-c). We found that all cell classes readily entrain to the sinusoidal ES, irrespective of ES amplitude and frequency (Figure 3a; N=24 pyramidal, 22 Pvalb, 13 SST cells). Specifically, ES induced a local V_e_ oscillation (measured from the closest extracellular electrode at 15 μm from the soma) that remained unaltered in amplitude (Figure 3b) and phase (Figure 3c) for 1-100 Hz. When looking at V_i_ and V_m_ (as estimated by the difference between V_i_ and V_e_), we found that the intracellular and the membrane potential at the soma of all cell classes followed all tested ES frequencies up to 100 Hz without any sign of membrane filtering (Figure 3b-c). Our findings are consistent with AC electrodynamics theory, predicting that the cell membrane voltage polarization induced by an external ES drive remains constant even for very high frequencies (e.g. ^38,39^). It follows that ES is a fundamentally different stimulus than intracellular stimulation where membrane RC filtering attenuates higher frequencies and allows lower frequencies to shape the membrane response. We also note that the amplitude of the pure ES effect induced was a fraction of 1 mV (V_m_: 0.29±0.26mV, Pyr; 0.36±0.23 mV, Pvalb; 0.34±0.11 mV, SST; Figure 3b). We conclude that all cells in the excitatory and inhibitory cell classes entrain to subthreshold ES unperturbed, from low to high frequencies (up to 100 Hz), without any apparent filtering, while the ES impact on the cellular membrane remains relatively modest in amplitude.

Is such unimpeded membrane entrainment across cell classes a result of the membrane residing at rest? To address this question, we depolarized and hyperpolarized all cells relative to their resting potential via intracellular DC injections and assessed the impact of ES on V_e_, V_i_ and V_m_ (Figure 3d-f). Simultaneous to the ES delivery we also delivered intracellular current steps (from −90 pA to 60 pA, duration: 5 s) to induce subthreshold-level changes in membrane polarization. The concurrent ES was set at 100 nA and 8 Hz. Once again, the subthreshold, ES-induced effect remained robust and unchanged across cell classes in terms of the V_i_ and V_m_-amplitude and -phase, even when the membrane polarization was varied by 40-50 mV (Figure 3d-f). In summary, we find that at subthreshold, rest and polarized levels, the cell membrane can follow the applied ES and does so robustly across all cell classes.

### Spike-field entrainment strongly corelates with the spike frequency of the cell

While cortical classes show strong, class-specific spike entrainment to ES, the same neurons exhibit indistinguishable entrainment characteristics to the same sinusoidal ES when not spiking: pyramidal neurons exhibit a combination of non-specific and frequency-specific (8 and 140 Hz) spike-field entrainment, inhibitory Pvalb neurons only entrain to high frequencies (140 Hz) while inhibitory SST entrain to medium-to-high frequencies (> 30 Hz). How can these observations be reconciled? To investigate whether cellular properties can explain ES entrainment, we considered four intrinsic properties – spike threshold, firing rate, input resistance and resting voltage ^36^. Looking at the intrinsic properties vs. the resulting spike-field strength (expressed by the vector length) we found that these properties do not explain within-class ES entrainment (Figure S7). With respect to between-class differences, intrinsic properties do not contribute to ES entrainment (see phase-distributions of control experiments in Figure 2). Therefore, ES is paramount for the emergence of spike-field entrainment even when cell classes differ in some of their intrinsic properties. We conclude that the stereotypical intrinsic properties do not explain the strong frequency-dependence seen in the spike-field coordination.

We next looked at the temporal structure of spiking for mechanisms that explain the ES frequency-dependence. To do so, we calculated the interspike interval (ISI; Figure 4a). Consistent with other studies ^36^, the instantaneous firing rate (for each spike, as calculated from 1/ISI) distributions show the stereotypical slow and irregular firing pyramidal neurons are known for (spike rate: 8.5±3.0 Hz; Figure 4b) as well as the fast, more stereotyped firing of Pvalb (133.4±35.7 Hz; Figure 4b). Finally, SST present an intermediate and more diverse case with faster spike rates than Pyr though slower than Pvalb and with a considerably wider firing rate-distribution (57.4±37.8 Hz; Figure 4b). Closer examination of the entrainment analysis suggests that at least a portion of the highly entrained spikes stem from neurons spiking at similar frequencies to the ES frequency, implying that for a given spike j, when (ES frequency) ≈ 1/ISI_j, the coordination between sinusoidal ES and spike j is enhanced. To test this hypothesis, we separated spikes based on whether their ISI was within a certain window of the ES frequency. For example, we looked at the spike-field coherence of pyramidal neurons with a spike frequency (1/ISI) of 8 Hz (“center frequency”) ± (frequency window) when the ES frequency is also 8 Hz (Figure 4c). We then varied the frequency window width and looked at whether the resulting population vector length changed within each bin. If no measurable difference is observed in terms of the vector length, then entrainment occurs equally strong across spike rate bins with no preference (in terms of ISI). If, on the other hand, spikes whose inverse ISI is close to the ES frequency are strongly entrained compared to the rest, then this supports a frequency-dependent mechanism like resonant ionic conductances where subpopulations of neurons are candidates to be highly entrained to a particular ES frequency depending on their ISI. The results from pyramidal neurons exhibit evidence for both phenomena (Figure 4c-d). While a subset of spikes (with 1/ISI ≈ ES frequency) are strongly entrained to the ES (Figure 4d), there is still substantial entrainment across pyramidal neurons (compared to control) and especially to ES frequencies much higher than pyramidal 1/ISI.

**Figure 4.**
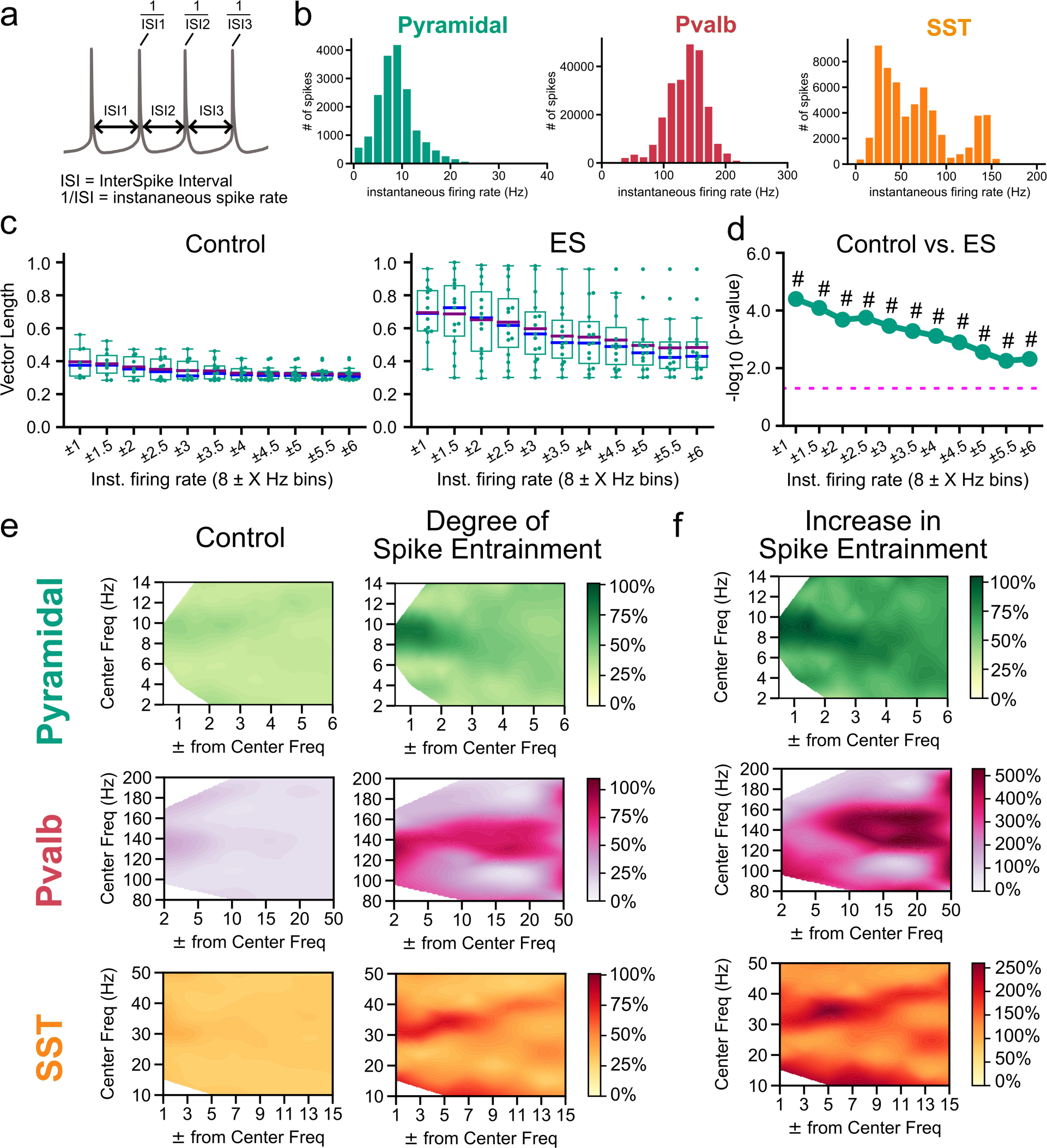
Cell class-specific ES entrainment of spiking correlates with spike rate properties of the individual classes. **a**, Schematic diagram demonstrating the instantaneous spike rate for each spike, calculated as the inverse of the interspike interval (ISI). ISI: time between a spike and the next consecutive spike. **b**, Histogram of the instantaneous spike rate distribution for all spikes recorded in pyramidal (green), Pvalb (red), and SST (orange) cell classes. (Pyramidal : N=21 cells, Pvalb: N=22 cells, SST: N=13). **c**, Degree of spike entrainment (as evaluated by population vector length) to ES (8 Hz and 200 nA) for each recorded cell’s spikes in the pyramidal class containing only spikes within a specific spike-rate range/bin (vector length means bootstrapped for 10000 trials). Bin size (boundaries) are designated by the spike rate to be analyzed (a “Center” Frequency) within a frequency-window range (ranging from ±1 to ±6 Hz). Tighter spike-frequency ranges are on the left, with the range widening towards the right. Boxplots: quartiles; purple and blue lines: mean and median values, respectively; whiskers: remaining distribution. **d,** Plot of -log_10_ p-values (Welch’s t-test; null-hypothesis: the two populations have equal means) for comparison between the same-spike-range bins (e.g., 8±1 Hz Control vs 8±1 Hz ES bins) between Control and ES (in **c**)). Dashed pink line: *p*=0.05; #: effect size (Cohen’s d) where d>0.8. **e,** Degree of spike entrainment (expressed as % of the normalized vector length, 0-1) to Control (no ES; left column) or ES application for spikes within specific spike-rate bins, in pyramidal (green), Pvalb (red), and SST (orange) classes (right column). Results shown for ES (200 nA) at 8 Hz for Pyr, 140 Hz for Pvalb, and 30 Hz for SST. **f**, Percentual increase in spike entrainment (vector length) in ES vs. Control for Pyr, Pvalb, and SST. Pyramidal: N=21 cells for ES frequencies 8, 30 Hz and N=13 for 140 Hz; Pvalb: N=22 cells for ES frequencies 8, 30 Hz and N=12 for 140 Hz; SST: N=13 for ES frequencies 8, 30 Hz and N=10 for 140 Hz.

To what extent do such highly entrained subpopulations of spikes exist in other cell classes? We used the same principle to look at spike-field entrainment as function of 1/ISI by varying the window (width) around the center frequency (see also Methods). The main difference between Pvalb and SST compared to pyramidal neurons is their discrepancy in spike rate (Figure 4b). For Pvalb, the condition 1/ISI ≈ ES frequency can only be met for ES frequency 140 Hz since Pvalb typically spike around 100-200 Hz. For SST the same condition holds for ES frequency around 30 Hz since they tend to spike around 20-100 Hz (Figure 4b). Spike-field entrainment of spikes at various ES frequencies is significantly increased when 1/ISI ≈ ES frequency across cell classes (Figure 4d-e). While this principle holds in general, the specifics of the entrainment profile differ between cell classes. For example, for pyramidal neurons at 8 Hz, spikes where 1/ISI deviates from the ES frequency by approx. 20%, exhibit a decrease in their degree of entrainment (vector length) by more than 50% (Figure 4e). On the other hand, for Pvalb at 140 Hz, 1/ISI can deviate by more than 35% (1/ISI≈90 Hz) without spikes experiencing an appreciable decrease in their entrainment. Finally, SST at 30 Hz form an intermediate case where spikes off by approx. 33% (1/ISI≈21 Hz) still exhibit 50-75% of the entrainment (Figure 4e). These observations are compatible with what is known about the cell classes explored in our study. Despite the entrainment of pyramidal neurons to subthreshold stimuli across ES frequencies, their irregular spiking due to a plethora of active and co-activated membrane mechanisms is a constant source of variability counteracting a tight entrainment window around the center frequency. On the other hand, once Pvalb with their fast and less adaptive spike-related mechanisms are entrained, there is little source of variability to counteract field entrainment which, in turn, results in a wide entrainment window around the center frequency. We conclude that class-specific spike-field coordination is the result of two effects: first, a non-specific membrane polarization ubiquitously present across cells and classes; second, class-specific excitability and activity properties that allow spikes of similar time scale to the ES to entrain particularly strongly. The latter effect is class-specific since how steeply spike-field entrainment drops for spikes where 1/ISI deviates from ES frequency varies across classes.

### Spike-field entrainment is dictated by the balance between membrane conductances but not by a specific conductance

The analyses pursued so far about the origin of the spike-field coupling point to a generic mechanism that cannot be explained only by a specific ion channel or morphological feature. To recapitulate, excitatory pyramidal neurons with their typically slower spike rate entrain to slow and fast ES, while inhibitory classes like SST and Pvalb with their fast spiking predominantly phase lock to fast ES. Importantly, the basic effect is replicated across regimes (subthreshold and spiking), cell classes (excitatory and inhibitory) and brain regions (mouse V1 and CA1). Even within the same brain area, the three cell classes we investigated have a different biophysical setup to each other, different expression profiles in many key ion channel genes (e.g., Kv3.1, HCN1-3, etc.) and different morphologies (pyramidal neurons are extended, Pvalb are more compact, SST are in-between).

We sought to test the hypothesis that the class-specific combination of conductances dictates the spike rate and, in this manner, the spike-field coupling to specific ES frequencies as opposed to a particular ionic conductance or morphology feature. To do so, we used biophysically realistic, computational single-cell models of excitatory and inhibitory V1 classes. Briefly, we recently developed a multi-objective single-cell generation workflow using whole-cell patch-clamp recordings (responding to step currents) and morphological reconstructions to generate and validate class-specific single-cell models^40^. Importantly, the evolutionary optimization procedure results in 40 best single-cell models per experiment termed “hall of fame” (or hof; Figure S8). The fact that a population of models is produced for each acquired cell (as opposed to a single “best” model) that generalize equally well offers the opportunity to assess mechanistic details as well as robustness – the same perturbation can be tested both across multiple models representing the same experiment/cell (“within cell”), across cells/experiments representative of a class (“within class”) and between cell populations that are members of different classes (“across class”).

We used bio-realistic single-cell models of pyramidal and Pvalb neurons and emulated the experimental setup by simulating the extracellular ES using a point-source approximation (Figure 5). As in experiments, we inject intracellular step currents combined with sinusoidal ES of various frequencies (8, 30 and 140 Hz; Figure 5a). The intracellular stimulation elicits spiking, with an overall spike rate broadly unaffected by simultaneous ES (Figure S9), but a spike phase distribution strongly affected by the presence vs. absence (control) of ES across frequencies for both pyramidal and Pvalb models (Figure 5a: control vs. ES rose plots for three ES frequencies). Specifically, the pyramidal model with a mean spike rate of approx. 8 Hz is entrained across ES frequencies (Figure 5a) whereas the Pvalb model with a mean spike rate larger than 100 Hz is only entrained for an ES frequency of 140 Hz (Figure S9). Therefore, the computational models used here capture the basic properties of spike-phase coupling observed in our experiments. We also tested how the spike frequency of a particular cell model affects spike-phase coupling to ES. To do so, we used the population hof models for the pyramidal and Pvalb cells and swept across the intracellular current step amplitude (pyramidal models, 180-250 pA; Pvalb models, 480-630 pA) keeping the resulting spike rate within a physiological range (pyramidal models, 1-14 Hz; Pvalb models, 100-160 Hz). As for the *in vitro* experiments, we saw that when the spike frequency of each hof model closely matches the ES frequency, the degree of spike-phase entrainment is maximized irrespective of the exact current injection value (Figure 5b, bottom row). Also like in experiments, the extent to which ES entrains spike phases varied across hof models (Figure 5b, bottom row).

**Figure 5.**
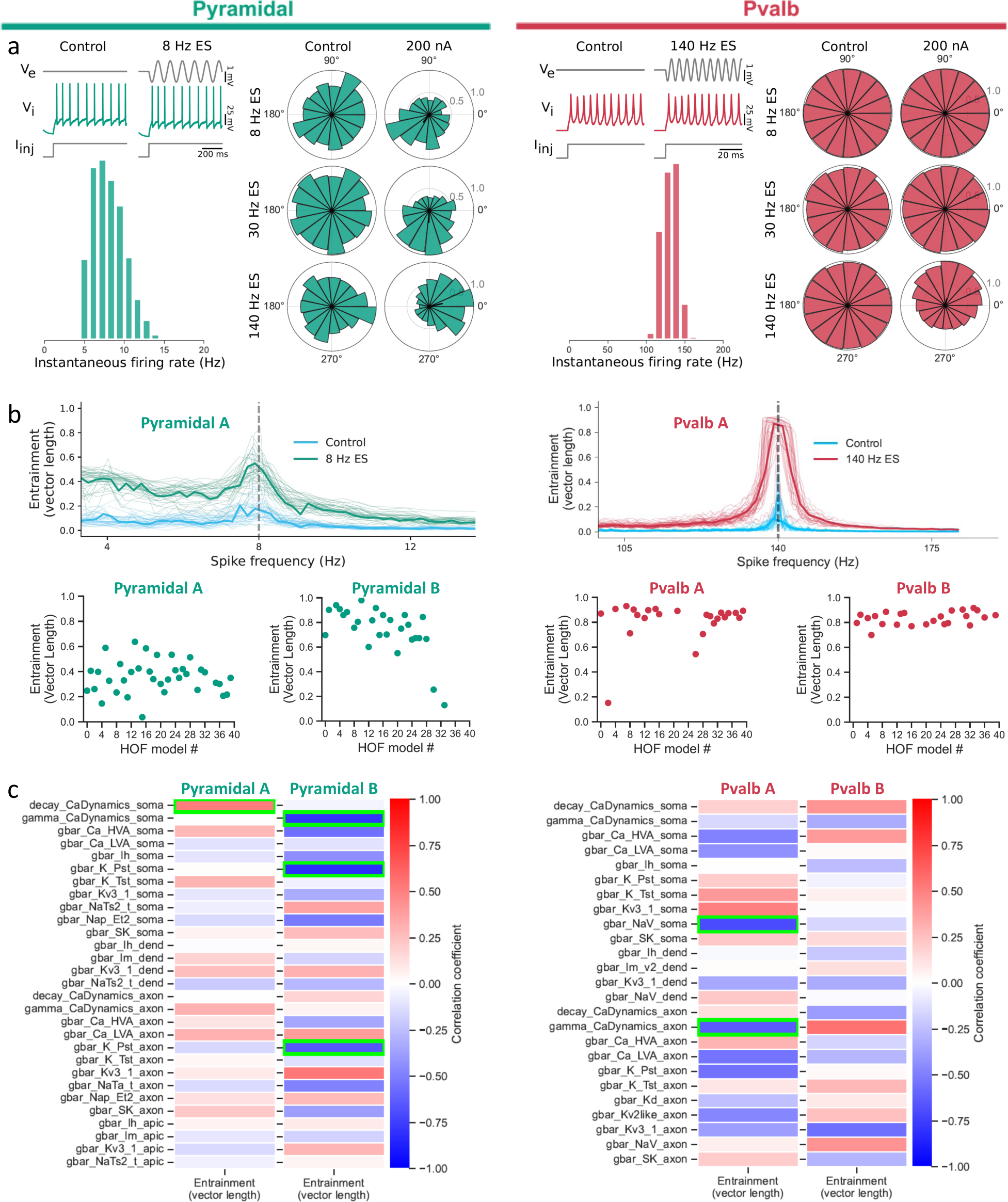
Computational modeling suggests spike rate differences rather than individual conductances as the major contributor to class-specific spike-field coupling. **a**, (Top left) A bio-realistic model of a pyramidal neuron (cell ID: 488698341) is used to emulate the experimental setup accounting for the sinusoidal ES. Intracellular DC current is combined with weak sinusoidal ES of various frequencies (8, 30 and 140 Hz) such that, just as in the experiments, the spike rate remains unperturbed by the ES). (Bottom left) Model ISI distribution (Control). (Right) Spike-phase relationship for the simulations (ES at 200 pA, top-to-bottom: 8, 30 and 140 Hz ES). Weak ES gives rise to strong spike-phase coupling in the presence but not in the absence of ES. (Top right) Same setup as for panel a but using a bio-realistic inhibitory Pvalb model (cell ID: 569998790; see also Figure S8). The Pvalb model shows preferential entrainment to fast ES while the pyramidal model readily entrains to both slow and fast ES (Figure S9). **b, Top,** Hall of fame (hof) models of the pyramidal (left)_and Pvalb (right) cell from panel a exhibit the robustness of the spike-field coupling (40 hof models per cell; Figure S8). Identical setup like in panel a for each hof model (spike-field entrainment: population vector length; thin lines: vector length for each hof model; thick line: mean vector length across hof models; cyan: control, no ES). With Pvalb spiking faster than pyramidal models (spike rate distributions in panel a), ES strongly entrains all hof models when ES frequency matches the spike rate. **b, Bottom,** Spike phase entrainment at the preferred ES frequency (pyramidal: 8 Hz; Pvalb: 140 Hz) across hof models for two pyramidal (pyramidal A: 488698341; pyramidal B: 354190013) and two Pvalb cells (Pvalb A: 569998790; Pvalb B, 471077857). **c,** Correlation between model conductances and spike-phase entrainment at the preferred ES frequency (for pyramidal neurons: 8 Hz; Pearson correlation across hof models; 40 hof models per cell; panel b, bottom row). Green boxes: statistically significant correlations (p<0.0017 for pyramidal and p<0.002 for Pvalb models). No conductance consistently correlates across the two pyramidal cells (i.e., compare locations of green boxes left vs. right column) despite the strong spike-phase coupling of individual hof models (panel b).

Do specific conductances dictate the spike-field coupling or does the combination of conductances dictate the spike rate that, in turn, affects entrainment to specific ES frequencies in a cell class-specific manner? To address this question, we looked at how individual conductances of the pyramidal and Pvalb hof models correlated with the degree of ES entrainment. For this analysis, we included a second pyramidal and Pvalb cell with their hof models (40 hof models per cell). We adjusted the intracellular current injection of each hof model to match the spike frequency with the ES frequency and found a distribution of spike-phase coupling strengths across hof models. To test whether specific conductances correlate with increased entrainment, we looked at the correlation between each hof model conductance and the spike-phase coupling (Figure 5c). For each cell, multiple conductances correlate with ES entrainment and exhibit statistical significance (Figure 5c, green boxes). But, when comparing models across the two pyramidal cells, very different sets of conductances correlate with vector length despite all hof models clearly showing strong spike-field entrainment at 8 Hz ES frequency (Figure 5b). For Pvalb, while all hof models clearly show strong spike-field coupling at 140 Hz ES frequency (Figure 5b) and several conductances correlate with entrainment across hof models of each cell (Figure 5c), yet no conductances consistently correlate with spike entrainment across the two Pvalb cells. We conclude that our simulations do not point to a specific conductance causing ES spike entrainment. Moreover, while these simulations do not completely exclude specific conductances as key contributors of spike-field entrainment, they suggest that the ES coupling in our experiments is not the result of a single conductance activated at a specific ES frequency but, instead, that the spike rate (i.e. the result of a balance between numerous conductances) of each cell class and its relationship to the ES frequency is a much stronger predictor of spike-field entrainment.

### Spike-field entrainment of human neocortical excitatory neurons to ES

To what extent do the ES effects seen in rodent V1 and validated in rodent hippocampal CA1 cells translate to human cells? While human pyramidal neurons share properties with their rodent counterparts (e.g. ^41,42^), they also possess many distinct characteristics (e.g. ^43,44^). The latter will impact their entrainment to ES. This, in turn, might affect how human vs. rodent neurons ought to be stimulated to achieve a certain activity pattern. We used our setup to investigate the subthreshold and spiking ES effects of human cortical (temporal or frontal lobe) neurons isolated from neurosurgical specimens ^45^.

We imposed the same, relatively weak sinusoidal ES field across human pyramidal neurons and measured how V_e_, V_i_ and V_m_ are affected by the ES strength and frequency (Figure 6a; field strength similar to rodent slices, compare Figure 6b to 1b) under two regimes spiking (Figure 6c-e) and subthreshold non-spiking (Figure 6f). For subthreshold ES, we found that human pyramidal cells entrain to subthreshold ES unperturbed, from low to high frequencies (up to 100 Hz), without any apparent filtering. Also, we found that at subthreshold, rest and polarized levels, the human cell membrane readily follows the applied ES across polarization levels robustly (Figure 6g).

**Figure 6.**
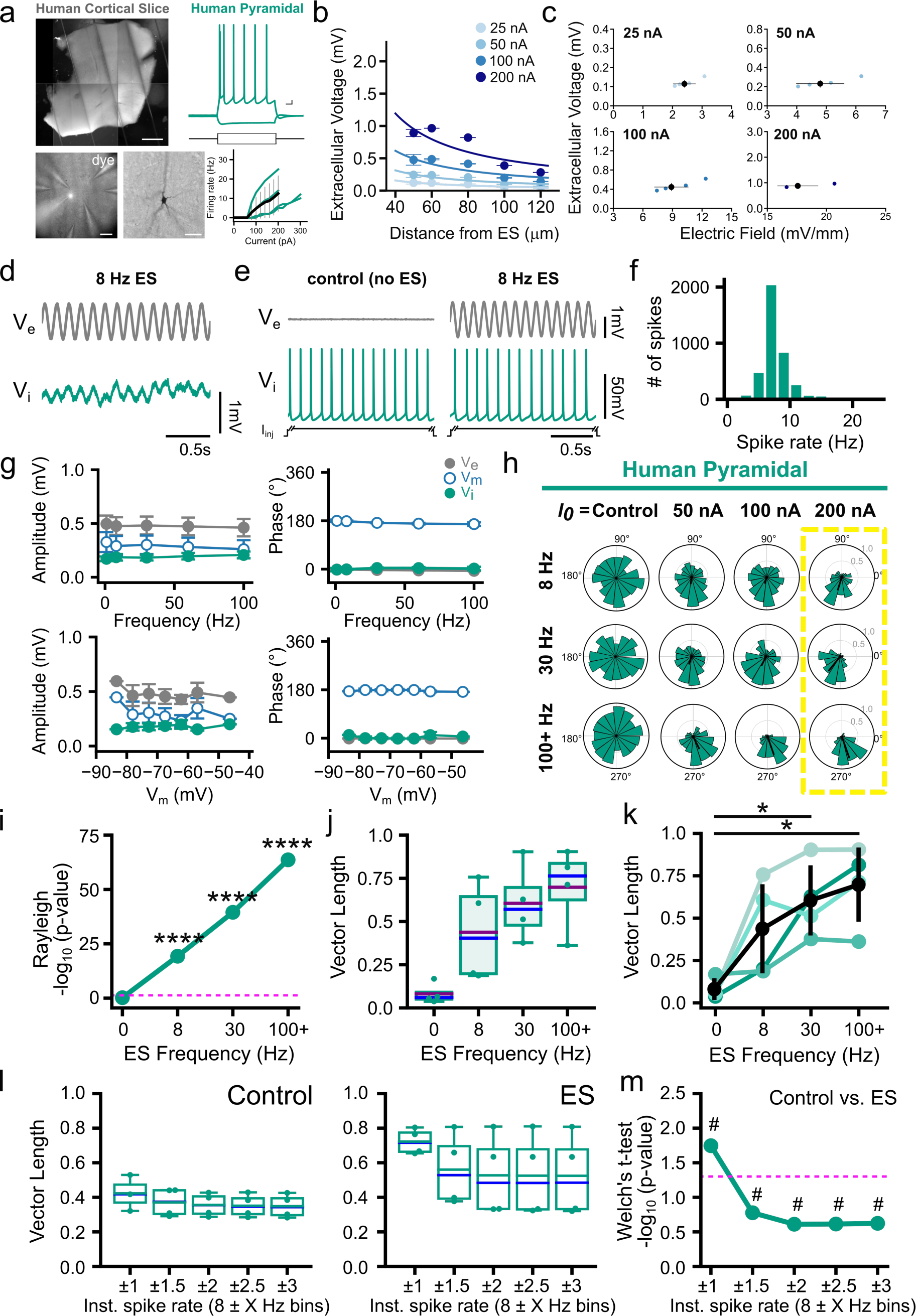
Sub- and spiking ES entrainment of human neurons. **a**, Top (left): Human *ex vivo* cortical slice obtained via neurosurgical tissue resections. (Right) Sample intracellular electrophysiology trace of human pyramidal neuron after depolarizing and hyperpolarizing current injections. Bottom (left): Human cortical slice with extracellular electrodes recording V_e_ at multiple locations 50-120 µm from pyramidal soma (white spot: soma of recorded cell filled with biocytin and dye for identification). (Middle): Pyramidal cell morphology of recorded cell visualized with post-hoc biocytin-HRP staining. (Right): f-I curves of recorded cells. **b,** Left: V_e_ amplitude as function of distance between ES and recording electrodes (ES amplitude: 25-200 pA; ES frequency: 8 Hz; circles: V_e_ mean amplitude; error bars: V_e_ amplitude st.d.) Trendlines: least-squares fit of the point source approximation. **c**, V_e_ and electric field amplitude elicited by the ES at the extracellular recording electrode closest to the whole-cell patched soma (approx. 15 µm). Blue: V_e_ amplitude for each experiment (n=5); Black: mean and st.d. across experiments. **d,** Sample trace of a human pyramidal neuron showing **e**ntrainment of V_i_ (green trace) to subthreshold ES. V_e_ (gray trace) measured at the closest extracellular location (15 µm) from the cell soma. The subthreshold sinusoidal ES is delivered 50 µm from cell soma (frequency: 8 Hz, amplitude: 100 nA). **e,** Simultaneous spiking induced by a suprathreshold DC stimulus I_inj_ that co-occurs with sinusoidal ES in human neurons. Despite its relatively small amplitude (see also **c**), the subthreshold ES affects neuronal spike timing without altering the spike number or frequency. **f,** Spike rate distribution for all spikes recorded from the human pyramidal neurons (N=4 cells). **g,** Subthreshold ES effect at resting potential (top) and at hyper- and depolarized potentials (bottom). Neurons were held at a range of membrane potentials via injection of depolarizing or hyperpolarizing current I_inj_ (from −90 to 90 pA). (ES amplitude: 100 nA, ES frequency: 1 to 100 Hz). V_e_ (gray)-, V_i_ (green)- and V_m_ (blue outlined circle)-amplitude (left) and phase (right) are shown (circles: mean; error bars: std). Human pyramidal neurons exhibit ES-frequency independence with induced V_i_-, V_e_- and V_m_-amplitude and -phase remaining constant across ES frequencies 1-100 Hz (top). The induced ES effect on V_i_, V_e_ and V_m_ amplitude and phase remain constant across a range of 40 mV polarization (bottom). **h,** Spike-phase distribution for varying ES parameters. (Rows) ES frequency (top to bottom): 8, 30, to 100+ Hz. (Columns) ES amplitude (left to right): 0 (Control), 50, 100 and 200 nA. Increasing ES amplitude increases spike-phase entrainment across slow (8 Hz), medium (30 Hz), and fast (100+ Hz) ES frequencies. **i-k,** Summary statistics for ES amplitude=200 nA (yellow box in h). **i,** Spike phase entrainment of human pyramidal neurons to ES assessed via Rayleigh’s test (****p ˂ 0.01; dashed pink line: *p*=0.05). **j**, The population vector length for each cell. Purple and blue lines: mean and median values, respectively; error bars: st.d. **k**, The population vector length for each cell (from **j,** each line represents a cell) across ES frequencies (paired t-test, false discovery rate (FDR)-corrected for multiple comparisons: *p<0.05, see Table S4). **l,** Spike entrainment (vector length) in Control (i.e. no ES, left) and ES (8 Hz and 200 nA) experiments (right) for each recorded human pyramidal cell, containing only spikes within a specific spike-rate range/bin. Bootstrap means of spikes within each cell obtained with 10000 trials. Bin size (boundaries) are designated by the spike rate (a “Center” Frequency) within a frequency-window range (ranging from ±1 to ±6 Hz). **m,** Spike phase entrainment assess via -log_10_ p-values (Welch’s t-test) for comparison between the same-spike-range bins (e.g. 8±1 Hz Control vs. 8±1 Hz ES bins) between Control vs. ES (in **k**). Dashed pink line: *p*=0.05; #: effect size (Cohen’s d) where d>0.8.

How about ES entrainment under spiking conditions? Like for rodent neurons, we injected intracellular step currents to elicit spiking simultaneously with the ES application. This did not affect their spike rate (paired *t*-test, p > 0.05 for control vs. 8 Hz, 30 Hz, and 140 Hz for all amplitudes tested) validating that, compared to the intracellular stimulation, the impact of ES on excitability [or number of spikes] is comparatively weak. Moreover, without ES (control), spikes are not entrained at the frequencies related to ES (Rayleigh testing, *p*>0.1; Figure 6g-k). Spike entrainment depends on ES and increases with ES amplitude (Figure 6h-k). Unlike their murine counterparts, human pyramidal neurons exhibit the strongest spike-field entrainment for high ES frequencies (140 Hz) despite their slower spiking (spike rate: 7.5±0.8 Hz). We also looked at the extent to which spikes with 1/ISI ≈ (ES frequency) entrain more strongly than the rest (Figure 6l-m). Given the slow spiking of human pyramidal neurons, this condition can only be satisfied for 8 Hz. We found that for 8 Hz, an ES frequency that human neurons entrain readily to, spikes with 1/ISI≈8 Hz entrain particularly strongly compared to spikes with a more disparate spike frequency, with inclusion of latter spikes resulting in the quick deterioration of coordination (Figure 6l-m). Thus, there is a subset of highly coordinated spikes where 1/ISI≈8 Hz. On the other hand, the very high entrainment of human excitatory spiking with fast ES frequencies, much faster than pyramidal cell firing, and the subthreshold entrainment properties of their membrane to fast stimuli also point to a mechanism very similar to the non-specific membrane polarization observed in rodent neurons. Similar trends are also observed even for weaker ES amplitudes (Figure S10). We conclude that, similar as in rodent neurons, spike-field coordination in human excitatory neurons is a combination of two effects: non-specific membrane polarization ubiquitously present across cells and a spike activity-related effect that allows spikes of similar time scale to the ES to entrain particularly strongly.

## Discussion

We found that membrane entrainment to a relatively weak (in amplitude) sinusoidal extracellular stimulus (ES) occurs across all cell classes (excitatory and inhibitory), multiple cortical areas (V1 and hippocampus) and species (rodent and human), pointing to a fundamental property of individual neurons. While intracellular current injections generally result in strong membrane filtering (i.e., higher frequencies are greatly attenuated by the membrane capacitance), membrane response to ES remains essentially unfiltered even for frequencies where intracellular stimuli are filtered out by the membrane capacitance (for frequencies >100 Hz ^46^). This is a generic property shared across all neurons and classes in our study. Yet, this generic property is superimposed onto a rich and diverse class-specific spike-field coordination. Specifically, we found that pyramidal neurons with their typically slower spike rate entrain their spiking to both slow and fast (up to 140 Hz) ES while inhibitory classes like Pvalb and SST with their comparatively fast spiking predominantly phase-lock to fast but not to slow ES. A resonant mechanism that purely relies on a match between spike rate and ES frequency may explain pyramidal coupling to 8 Hz ES as well as Pvalb coupling to 140 Hz but cannot explain the pyramidal and, to a lesser extent, SST coupling to very fast ES. We note that the class-specific spike-field coordination stems from the use of sinusoidal ES rather than alternative waveforms (e.g., pulses) where the fundamental frequency of the stimulus is obscured, and the response occurs at multiple time scales. We conclude that sinusoidal ES entrains subthreshold and spiking activity across cell classes, resulting in a rich repertoire of suprathreshold responses.

Our experiments point to class-specific spike-field coordination resulting from two effects. The first one is a non-specific membrane polarization ubiquitously present across all cells. It is exemplified in the subthreshold ES experiments where membrane ES entrainment is universal and does not depend on the specifics of the type or area. The second effect is class-specific with spikes elicited at similar time scales to the ES strongly phase-locked to the external field. While some ionic conductances have been linked to resonant coupling at different frequencies like the m-current ^47,48^ and the hyperpolarization-activated cation h-current^42,49^, the spike output response of neurons is ultimately the result of interactions between multiple ion channels and their morphologies. Our experiments and computational models show that distinct cortical classes with different conductance and spiking profiles exhibit ubiquitous and robust coordination to ES across frequencies. This suggests that membrane entrainment by the ES is a fundamental property rather than mediated by a specific ion conductance (though the presence of specific conductances can increase or decrease coupling properties, e.g.^47,48^). Still, since the cellular response (from the spike waveform to the excitability profile and the resulting spike rate) is crucially affected by its ionic setup, a cell’s active properties are implicitly reflected in the ES entrainment. For example, while V1 pyramidal neurons typically spike slowly, their spike rate can increase drastically during bursting (i.e., 200-300 Hz), mediated by a host of soma-dendritic mechanisms, suggesting that their ionic setup, in principle, supports fast activity and locking to fast ES ^50–55^. We conclude that while ES entrainment is ubiquitous, the characteristics of spike-field entrainment for each cortical class ultimately reflects its morpho-electric properties and ionic setup.

Our results in V1 and CA1 layer 5 pyramidal neurons of adult mice bear some similarities with the observations made for somatosensory layer 5 pyramidal neurons in the juvenile rat^19^: subthreshold sinusoidal ES coupling is ubiquitous, unfiltered (even at ES frequencies up to 100 Hz) and robust while small-amplitude ES is very potent at biasing the spike phase (but not spike frequency). Also, both juvenile and adult pyramidal neurons exhibit preferred spike coupling to 8 Hz ES vs. 30 Hz ES when compared to control. But here we also report how fast ES beyond 100 Hz leads to strong spike-field coupling in pyramidal neurons, a surprising observation given the slower (compared to the ES frequency) spike rate of pyramidal neurons in our study. It is conceivable that pyramidal neurons in juvenile rat vs. adult mice may entrain differently to fast ES due to age-related differential expression of individual ion channels impacting the spike response even if the subthreshold response remains similar ^19^.

In contrast to rodent excitatory neurons, human layer 5 pyramidal neurons entrained across ES frequencies with particularly strong phase-locking at fast ES (140 Hz). Where does such strong spike-field coordination come from? Notably, the subthreshold response of human pyramids is not different to mouse V1 or CA1 neurons so that their increased entrainment for fast ES cannot be solely explained from the subthreshold results. It has been suggested that human neurons have lower capacitance filtering compared to rodent pyramidal neurons^43^. Decreased filtering may contribute to strong spike coupling for fast ES as well as enhanced integration of fast synaptic input ^56,57^. The hyperpolarization-activated non-specific cation current Ih, exhibits higher expression in human vs. mouse neocortical excitatory neurons with increased Ih expression counteracting dendritic filtration^42^. Finally, the voltage-gated potassium conductance at the soma of layer 5 human pyramidal neurons is smaller (normalized for size) than in rodents and can contribute to spike repolarization and timing ^58–60^. While our computational models did not identify a specific role for these conductances in rodent neurons, they may contribute in spike-field coupling of human neurons and their preferential phase-locking at fast ES (140 Hz).

Our findings shed light on cellular ES effects and are relevant for the design of next-generation neuromodulation technologies using customized ES waveforms for the selective coordination of specific cell classes *in vivo*. The main parameter exerting such selectivity based on our results is the ES frequency, with slow ES facilitating coordination of excitatory classes while fast ES facilitates coordination of both excitatory and inhibitory populations. Spike-field coordination in various frequency bands is the substrate of many cognitive functions and therapeutic interventions ^34,61–63^. Beyond local ES effects in the vicinity of the stimulation site, class-specific ES protocols can impact spatially distributed brain regions away from the stimulation site. For example, in humans, cortical structures have reliable monosynaptic connections to the subthalamic nucleus (STN), a major target for deep brain stimulation in movement disorders ^4,6,64–68^. Class-specific stimulation of cortex via ES may allow the selective neuromodulation of individual STN classes based on their projection pattern and, in such manner, alter behavior or facilitate therapy. Beyond invasive intracranial stimulation administration, we expect the same effects to apply to noninvasive stimulation as long as they impose an electric field of certain spatiotemporal properties across the membrane of neurons (e.g.^69^). Our findings form the basis for designing such selective, class-specific ES interventions.

Coordinated circuit activity often produces an endogenous and measurable extracellular field, especially so in cortical structures like the hippocampus and neocortex^70^. In addition, field potentials can also be detected distant from original generator site, e.g., hippocampal theta oscillations can be volume-conducted and measured in adjacent neocortical regions^71^. Because the field strength in our experiments (∼1-25 mV/mm) is comparable to what is measured *in vivo*, our results can also shed light in cellular entrainment due to endogenous fields, *i.e.,* via ephaptic coupling ^72^ or far field (i.e. the location in space in which the field can be approximated by a dipole) effects from other regions with the spatial coherence and geometry of the sources impacting the spatial extent of the field^73^. The ubiquitous nature of sinusoidal ES coupling suggests that slower (e.g., alpha, theta, beta) and faster endogenous oscillations (gamma, ripples) are felt and followed by the membrane across cortical neurons. To what extent is spiking affected by such oscillations in a manner that constitutes an alternative, non-synaptic communication between neurons? In the absence of synaptic drive that sufficiently depolarizes neurons and leads to spiking, endogenous fields are rarely strong enough to trigger spiking under physiological conditions. But, in the presence of such synaptic drive (that often results in an extracellular field, ^70^), electric fields can modulate spike timing and enhance the coordination between local spiking and various bands of the local field potential. For example, Purkinje cells in the cerebellum coordinate their spiking with ms precision through ephaptic coupling ^26^. Fast coordination (within ms) between pyramidal neurons has also been observed in neocortex (e.g. ^53^) despite local synapses only supporting slower spike coordination. The spike-field coupling we see across classes for fast ES (140 Hz) can explain such fast coordination even in the absence of very fast synapses and, in such manner, provide an alternative, energy-efficient, non-synaptic paradigm for circuit coordination across timescales in cortical structures.

## Acknowledgments

We thank the Allen Institute Transgenic Colony Management, Tissue Processing, Histology, Imaging, Morphology, and 3D reconstruction teams for their assistance in this work, and the surgeons at the University of Washington Medical Center, Swedish Medical Center and Harborview Medical Center for human tissue collection. We thank Ueli Rutishauser for discussions. CAA thanks the Board of Governors of Cedars-Sinai Medical Center. Research reported in this publication was supported by the National Institute Of Neurological Disorders And Stroke of the National Institutes of Health under Award Numbers R01NS120300 and RO1NS130126. The content is solely the responsibility of the authors and does not necessarily represent the official views of the National Institutes of Health. We thank the Allen Institute founder, Paul G. Allen, for his vision, encouragement, and support.

## Contributions

Conceptualization: CAA; Methodology: SYL, KK, FB, LC, TJ, CK, CAA; Investigation: SYL, KK, CAA; Visualization: SYL, KK, CAA; Funding acquisition: SYL, CAA; Project administration: SYL, CAA; Supervision: CK, CAA; Writing – original draft: SYL, CAA; Writing – review & editing: SYL, KK, FB, LC, TJ, CK, CAA.

