## supplemental material for "Cell class-specific electric field entrainment of neural activity"

**Summary:** Electric fields affect the activity of neurons and brain circuits, yet how this interaction happens at the cellular level remains enigmatic. Lack of understanding on how to stimulate the human brain to promote or suppress specific activity patterns significantly limits basic research and clinical applications. Here we study how electric fields impact the subthreshold and spiking properties of major cortical neuronal classes. We find that cortical neurons in rodent neocortex and hippocampus as well as human cortex exhibit strong and cell class-dependent entrainment that depends on the stimulation frequency. Excitatory pyramidal neurons with their typically slower spike rate entrain to slow and fast electric fields, while inhibitory classes like Pvalb and SST with their fast spiking predominantly phase lock to fast fields. We show this spike-field entrainment is the result of two effects: non-specific membrane polarization occurring across classes and class-specific excitability properties. Importantly, these properties of spike-field and class-specific entrainment are present in cells across cortical areas and species (mouse and human). These findings open the door to the design of selective and class-specific neuromodulation technologies.

### METHODS AND SUPPORTING MATERIAL

#### Methods

Acute Slice preparation:

Mouse and human tissue were prepared with slice preparation reagents and protocols standardized at the Allen Institute [74].

Acute mouse brain slices, from both female and male mice at ages P40-P50, were prepared as described in [75]. All mouse procedures were approved by the Allen Institute's Institutional Care and Use Committee. Mice were anesthesized with 5% isoflourane and transcardially perfused with ice-cold, oxygenated artificial cerebrospinal fluid solution (ACSF.I[74]): in mM: 0.5 $CaCl_2 \cdot 2H_2O$, 25 D-glucose, 98 HCl, 20 4-(2-hydroxyethyl)-1-piperazineethanesulfonic acid (HEPES), 10 $MgSO_4 \cdot 7H_2O$, 1.25 $NaH_2PO_4$, 3 myo inositol, 12 N-acetylcysteine, 96 N-methyl-D-glucamine (NMDG), 2.5 KCl, 25 $NaHCO_3$, 5 Na-L-ascorbate, 3 Na-pyruvate, 0.01 taurine, 2 thiourea). Parasagittal cortical slices (350 µm) were prepared in ACSF.I with a Compresstome (Precisionary Instruments) or VT1200S Vibratome (Leica Biosystems), and the slicing angle was set to 17 degrees relative to the sagittal plane to preserve apical dendrites of pyramidal cells. Hippocampal slices were sliced in the coronal plane. Slices were kept in a recovery chamber containing oxygenated ACSF.I for 10 minutes at 34ºC. They were then transferred to a holding chamber containing oxygenated ACSF.IV[74], consisting of the following (in mM): 2 $CaCl_2 \cdot 2H_2O$, 25 D-glucose, 20 HEPES, 2 $MgSO_4 \cdot 7H_2O$, 1.25 $NaH_2PO_4$, 3 myo inositol, 12.3 N-acetyl-L-cysteine, 2.5 KCl, 25 $NaHCO_3$, 94 mM NaCl, 5 Na-L-ascorbate, 3 mM Na-pyruvate, 0.01 taurine, 2 thiourea. Slices were kept in the holding chamber (at least one hour) at room temperature until time of recording.

Acute human cortical slices were prepared as described in [75]. All procedures involving human tissue are reviewed by the review boards at the hospitals performing the surgeries before beginning the study, and all patients provided informed consent. Tissue was extracted from the temporal or frontal lobe of patients undergoing neurosurgery for epilepsy or tumors (mean age: 27 ± 13.8 years, 3 donors). The tissue specimens were distal to the core pathological site and was deemed not to be of diagnostic value. The specimens were placed in a sterile container filled with pre-chilled, carbogenated aCSF VII[74] (in mM: 0.5 $CaCl_2 \cdot 2H_2O$, 25 D-glucose, 92 HCl, 20 HEPES, 10 $MgSO_4 \cdot 7H_2O$, 1.2 $NaH_2PO_4$, 92 NMDG, 2.5 KCl, 30 $NaHCO_3$, 5 Na-L-ascorbate, 3 mM Na-pyruvate, 2 thiourea). Specimens were transported from the surgical site to the laboratory within 10-40 min, where they were trimmed and mounted for best preserving intact cortical columns (pial surface to white matter) during slicing. Specimen were sliced in aCSF VII using a Compresstome or Vibratome and transferred to a holding chamber containing oxygenated aCSF VII at 34°C for 10 min, then moved and kept in aCSF IV[74] (REF) at room temperature for a minimum of one hour before recording began.

Electrophysiology and analysis

Rig setup:

Recordings were performed in a customized multi-patch system, as described in [75]. The multi-patch system consists of 8 headstages with pipettes arranged in a semi-circular organization, as seen in Figure 1a, and allows for simultaneous intra- and extracellular stimulation and recordings at multiple locations close by cell soma. One pipette was used for the intracellular stimulation and recordings, one for extracellular stimulation, and the rest were placed within 50-120 $\mu m^3$ of each cell soma to record the extracellular voltage. The electrode distance and placement (within +/- 5mm) was measured with Acq4 software [76], which was connected to the microscope and micromanipulators for providing accurate tracking of the hardware and cell position. The pipette holders were equipped with customized metal shields to reduce crosstalk artifacts. Each headstage was controlled independently through modified triple-axis motors (Scientifica; PatchStar). Intracellular and extracellular stimulation was applied through Multiclamp 700B amplifiers (Molecular Devices). (Intracellular and extracellular stimulation was applied through different amplifers). Data acquisition was conducted using Multi-channel Igor Electrophysiology Suite (MIES; https://github.com/AllenInstitute/MIES), custom software written in Igor Pro (WaveMetrics). Recorded signals were amplified with a Multiclamp 700B amplifier and digitized at 50-200 kHz with ITC 1600 DAQs. (Heka). Pipette pressure was regulated using both electro-pneumatic control valves (Proportion-Air; PA2193) under control of MIES software, and through manual (via mouth) pressure. Slices were visualized through an upright microscope (BX61WI, Olympus) with oblique infrared illumination (WI-OBCD condenser) equipped with 4X and 40X objectives, on a custom motorized stage (Scientifica), and images were taken with a digital sCMOS camera (Hamamatsu; Flash 4.0 V2). Acq4 software[76] was also used for imaging, and subsequent image analysis.

Cell class identification and characterization

In mouse recordings, individual neuronal cell classes were identified by the following methods: Excitatory pyramidal cells were identified with Cre- or Flp-dependent transgenic mice with fluorescent reporters (TdTomato or eGFP) specific for Layer 5 excitatory neurons (through Tlx3 or Sim1 promoters) or by their pyramidal morphology (in V1 and hippocampus). They were confirmed by their regular-spiking firing pattern (in response to intracellularly injected current steps) in patch-clamp recordings. Inhibitory Pvalb and SST cells were identified with Cre- or Flp-dependent transgenic mice with fluorescent reporters (TdTomato or eGFP) specific for Pvalb or SST cell classes (though Pvalb or SST promoters), and confirmed by their fast-spiking firing pattern. The intracellular recording pipette was filled with dye (Cascade Blue) to confirm overlap between the recorded cell and fluorescence to ensure proper cell class targeting. In human neuron recordings, pyramidal classes in Layer 5 were targeted by their location (approximately 1800 mm from pia [77]), large size and clear apical dendrites, and then confirmed by their regular-spiking firing pattern (in response to intracellularly injected current steps) in patch-clamp recordings.

After recordings, the slices underwent *post-hoc* processing to confirm cortical layer boundaries (via 4',6-diamidino-2-phenylindole (DAPI) staining) and to reveal their cellular morphology (via biocytin labeling). Slices were first processed for immunohistochemistry by fixation in 4% paraformaldehyde and 2.5% glutaraldehyde for at least 40 hours at 4°C, then transferred and washed in phosphate buffer saline (PBS) for 1-7 days. Slices were then placed in 5 µM DAPI in PBS for 15 min at room temperature and then washed three times in PBS. Slices were then transferred to 1% hydrogen peroxide ($H_2O_2$) in PBS to extinguish endogenous peroxidases for 30 min, and then washed three times in PBS. For biocytin labeling, a 3,3'-diaminobenzidine (DAB) peroxidase substrate kit (Vector Laboratories) was used. Slices were mounted and dried for approximately 2 days before imaging on an AxioImager Z2 microscope (Zeiss) equipped with an Axiocam 506 monochrome camera (Zeiss) at 20X, with images acquired via the Zeiss Efficient Navigation software.

Computer simulations

Single cell simulations were carried out in NEURON (Version 8.2.2) [78], using the BMTK Python framework (BMTK Version 1.0.7 – Python Version 3.10.11)[79]. Bio-realistic, conductance-based mouse V1 single-cell models developed by Nandi et al.[40] were adopted. Briefly, we recently developed a workflow to generate biophysically and morphologically detailed cortical single-cell models at scale. It relies on a parallel multi-objective optimization framework deployable in high-performance computer architectures[40,80,81]. From patch-clamp experiments, electrophysiology responses to a battery of standardized current stimuli (1 s-long DC current injections of increasing amplitude) are analyzed, resulting in a set of subthreshold and spiking features for each experiment (e.g., spike timing, amplitude, width, etc.) The workflow extracts 11 electrophysiology features from *in vitro* experiments, and their mean and standard deviation (std) are computed for a particular stimulation waveform. For every feature, an absolute standard score is calculated $Z_i=|f_i-\mu_i|/\sigma_i$ with the current feature value ($f_i$) measured from the output traces of the models and $\mu_i$, $\sigma_i$ being the experimentally measured mean and std, respectively[82]. The second data modality required for the biophysically realistic single-cell models is the morphology (soma and dendritic compartments, axonal ones are not required but can also be used). Axonal reconstructions were replaced with a stub using the Import3d_SWC_read() NEURON function.

We replicated experiments by simulating 10 s-long segments and evaluated data from 1 to 9 s (for Pvalb cells) or by simulating 20s-long segments to evaluate data from 1 to 19 s (for Pyramidal cells), with intracellular current step injections of varying amplitude. The ES electrode was positioned 50 μm from the soma in agreement with the *in vitro* experiments. Intracellular and extracellular current injections were configured using the IClamp and Xstim BMTK modules respectively. Two cell classes (pyramidal and Pvalb) and two different cells from each class were simulated. Corresponding cell IDs: Pyramidal A, 488698341; Pyramidal B, 354190013; Pvalb A: 569998790; Pvalb B, 471077857. In addition to the intracellular DC injection, we also added continuous background noise (Gaussian distribution) with an amplitude optimized to replicate the experimental spike rate distribution statistics. Specifically for Figures 5a and S9a, multiple simulations of hof 0 for Pyramidal A and Pvalb A models were performed (Pyramidal: 16 simulations 194-240 pA; Pvalb: 48 simulations 488-630 pA; N = 2,453 spikes for each Pyramidal rose plot, N = 48,942 spikes for each Pvalb rose plot). For the simulations in Figures 5a,b, the std of the intracellular noise injection was 65 pA for the Pyramidal and 110 pA for the Pvalb models leading to a spike rate standard deviation of 1.5 Hz and 5 Hz, respectively. For the simulations in Figure 5c, we adjusted the injected DC stimulus and noise distribution, with the pyramidal models having a spike rate of 8±1.5 Hz (mean±std) and the Pvalb models having a spike of 140±5 Hz (mean±std). The target spike rate for the models sought to replicate the experimental spike rate across models of the same class (Table S1). The maximum accepted error offset was ± 0.1 Hz. Models which failed to satisfy these criteria were not used for the correlation analysis (Figure S9).

<u>Electrophysiological recordings</u>

Electrophysiological recordings were performed in slices (for both mouse and human slices) at 32-34 °C perfused under carbogenated recording ACSF (aCSF.IX[74] (in mM: 1.3 $CaCl_2 \cdot 2H_2O$, 25 D-glucose, 1 $MgSO_4 \cdot 7H_2O$, 1.2 $NaH_2PO_4$, 3 KCl, 18 $NaHCO_3$, 126 NaCl, 0.16 Na-L-ascorbate). To examine the effects of ES without potentially confounding factors of synaptic activity, recordings were conducted in the presence of synaptic blockers. Synaptic transmission blockers kynurenic acid (1 mM, Sigma) and gabazine (10 μM, Tocris) were added to the aCSF to block fast glutamatergic and GABAergic synaptic activity, respectively. Whole-cell patch-clamp recordings were performed with internal solution containing 130 K-gluconate, 10 HEPES, 0.3 ethylene glycol-bis(β-aminoethyl ether)-N,N,N',N'-tetraacetic acid (EGTA), 3 KCl, 0.23 $Na_2GTP$, 6.35 $Na_2$Phosphocreatine, 3.4 Mg-ATP, 13.4 Biocytin, and 50 μM Cascade Blue dye (excited at 490 nm), 280-295 mOsm and pH 7.2-7.3. (Electrophysiological values are reported without liquid junction potential correction). Pipettes for extracellular stimulation and voltage recordings contained aCSF IX. Patch pipettes were pulled from borosilicate glass capillaries (Sutter Instruments) with a DMZ Zeitz-Puller (Zeitz), with tip resistances of 4-6 MΩ for whole-cell recordings and 1-2 MΩ for extracellular stimulation and voltage recordings.

Whole-cell patch-clamp recordings were performed on neurons that had a stable seal (>1 GΩ) and successful break-in. Bridge balance was monitored and did not exceed 15 MΩ. Resting membrane potential (RMP) was measured at 1-2 minutes after break-in of the cell, and only neurons with RMP over -53 mV were recorded. Bias current was injected within the MIES software package for the remainder of the experiment to maintain the determined resting membrane potential. To examine the cell-class-specific firing pattern and intrinsic properties of the recorded cell, and to generate the F-I (frequency-current) curve, intracellular current steps were injected into the cell at current-clamp mode. 20 pA steps (1 second) from -160 pA up to 200 pA were applied to the cell for all cell classes), and in further steps up to 400 pA for cells with a higher spike threshold. The rheobase was determined by the minimum current amplitude needed to generate an action potential (AP). The interspike interval (ISI), the difference in time between an APl and the next subsequent AP, was used to calculate the cell firing rate. AP threshold was calculated by first identifying the membrane voltage deflection point at which the first-order derivative of the membrane potential (dV/dt) exceeded 20mV/ms, and adjusted to where the dV/dt was 5% of the average maximal dV/dt. Input resistance was calculated by first plotting the voltage vs. the injected current at the hyperpolarizing steps from -120 pA to 20 pA, and then taking the slope of the resulting fit.

Subthreshold entrainment and analysis

To analyze the effect of ES on subthreshold cellular dynamics, we recorded from neurons under current-clamp mode, while delivering ES of varying amplitude (25-200 nA) and frequency (1-100 Hz) for 5 seconds. Neurons were either held at their resting potential, or at varying subthreshold polarization levels (through intracellular current delivered through the pipette 30 pA steps, from -90 pA to 90 pA). To calculate the resulting amplitude, frequency and phase of the extracellular ($V_e$), intracellular somatic ($V_i$), and the resulting membrane voltage ($V_m$, calculated as $V_m$=$V_i$-$V_e$), we first divided the raw unfiltered trace into [0, T]-timed intervals, where T (in seconds) represents the period of the ES as defined by T = 1/frequency. (e.g. for an 8 Hz stimulus, T = 0.125 s (1/8 Hz), which results in 40 intervals (5 s / T) for a 5 second stimulus duration). We then aligned these intervals to determine the mean waveform by calculating the mean of the aligned interval traces from 0 to T. The $V_e$ trace used was from extracellular electrode closest to the cell soma (15 μm away), and the $V_i$ trace was from the intracellular recording electrode. The amplitude, frequency, and phase of the mean waveform of each neuron from each recording was extracted for the analysis of subthreshold neuronal entrainment in Figure 3.

Spike entrainment and analysis

To examine the effects of ES on the suprathreshold properties of neurons, we injected intracellular DC current into neurons under current-clamp mode, to elicit action potentials during a 9 second intracellular stimulation protocol. Neurons were held at their resting membrane potential, and the DC current was first injected into the neurons in the absence of ES ("Control" condition), and then same DC current was injected into the neurons while concurrently applying ES at varying frequencies (8-140 Hz) and amplitudes (25 to 200 nA). (DC current amplitude was determined as the current injection necessary to sustain spiking during the protocol) For the spike-phase calculation, i.e. the phase of the extracellular $V$e when an action potential is elicited, the $V_e$ trace was first bandpass-filtered with frequency limits from 0.5 Hz to 200 Hz. The $V_e$ trace used was from extracellular electrode closest to the cell soma (15 μm away). We then used the Hilbert transform to calculate the instantaneous phase of the oscillating $V_e$, closest to the cell soma during spiking. The phase (in degrees) was measured in the range 0° to 360°, with 90° the peak and 270° the trough of the $V_e$ waveform. Traces were analyzed after the first 0.5 s of the protocol to ensure stability of cell spiking for analysis. Spike time for each action potential was measured individually for each spike at the voltage deflection point at which the first-order derivative of the membrane potential (dV/dt) exceeded 20 mV/ms. The spike phase is thus defined as the instantaneous $V_e$ phase at the spike time. Spike times were also used to count and compare the number of spikes during Control vs. ES-application conditions.

We examined the spike phases during the ES via circular statistics (with the ES-modulated $V_e$ as reference) to determine whether a neuron was significantly phase locked. The Rayleigh test was used to assess the uniformity of spike-phase distribution (null hypothesis: distribution is uniform), and population-vector analysis was used to examine whether spikes exhibit a preference to any specific phase of the imposed $V_e$. The spike distribution during control (i.e. no ES) experiments was examined during alignment to a "virtual" (using the $V_e$ from the subsequent ES experiment) extracellular field and compared to the spike distribution of neurons during ES.

The relationship between spike rate and spike-field entrainment was examined in detail by taking the vector lengths of spike dataset per cell, and binning the spikes according to their instantaneous spike frequency (as calculated by the spike ISI). Bootstrap mean values for each cell in each cell class was obtained by shuffling the number of spikes in 10000 trials. The spike-frequency bins were constructed by taking the specific ES frequency inducing entrainment for each cell class (a "center frequency") and adjusting the bin width by adding/subtracting from the center frequency. The degree of entrainment (as assessed by vector length) for the specific spike-frequency bins, was determined by taking the values of the spike-frequency bins (during ES) and dividing them by the values of the spike-frequency bins during Control conditions.

Statistical analyses

Statistical analysis was performed was performed using Python in SciPy[83] and Circstat (https://github.com/circstat/pycircstat). Data was tested for normality (Shapiro-Wilk test), and for normally distributed data, we used the two-sided Welch's t-test or paired t-test. One-way ANOVAs were used to compare multiple frequencies in Figure 2, and the Cohen's test and Levene's test were used to examine effect size and homogeneity of variance, respectively, as indicated in the figures. Significance was determined at $p<0.05$, as denoted by the following: (* $p < 0.05$, **: $p < 0.01$, *** $p < 0.001$, **** $p < 0.0001$). Statistical tests performed with multiple comparisons are reported with false discovery rate (FDR)-corrected p-values using the Benjamini-Hochberg method. Values in the text are presented as mean ± standard deviation, or where as noted, also with median and interquartile ranges.

**Figure S1**

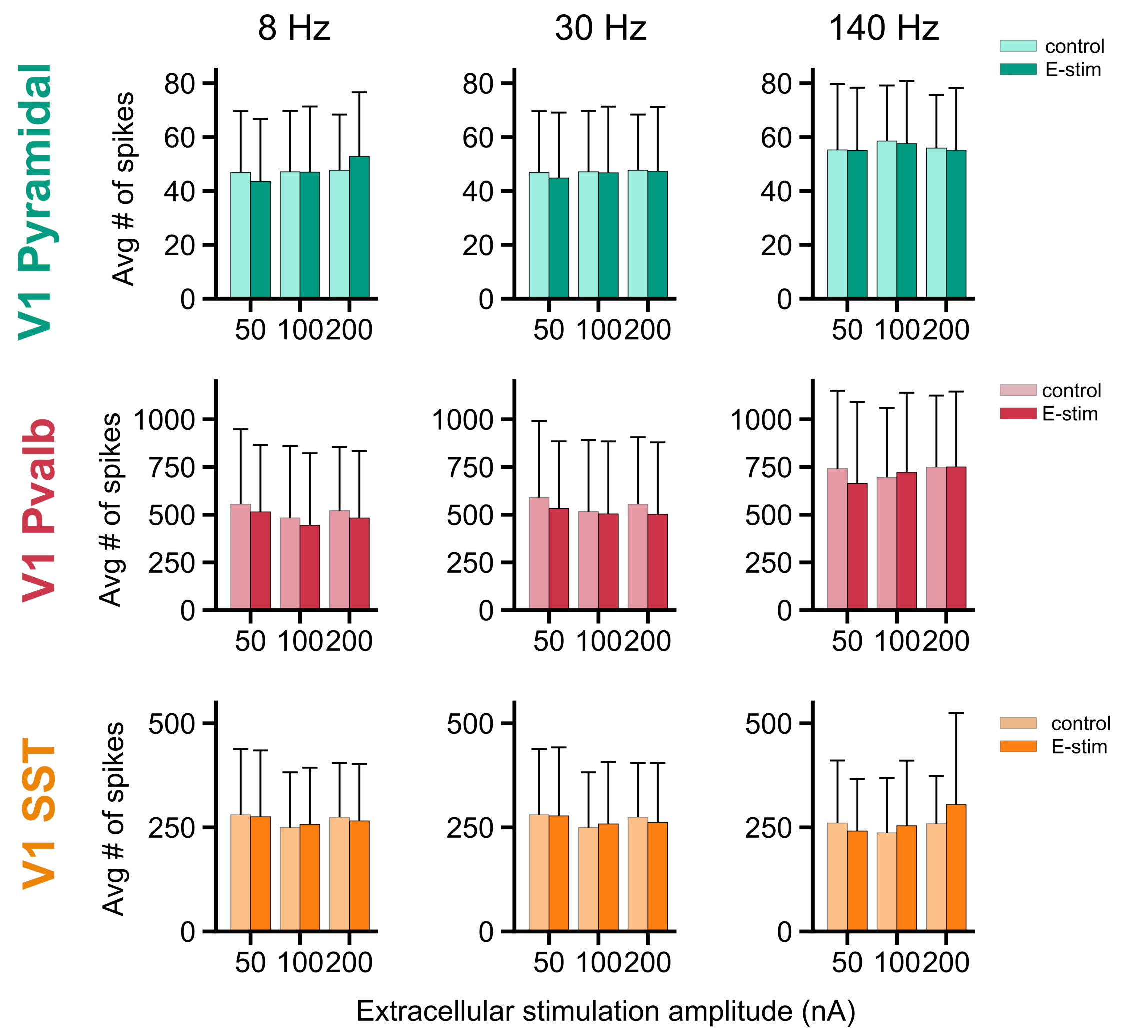


**Figure S1. Extracellular ES does not affect spike rates across classes.** To assess whether ES affects the spike rate, we examined the average number of spikes during both control (no ES) and ES. Plots are shown comparing the average number of spikes evoked during control vs. the varying ES conditions (in amplitude, frequency), for each cell class. None of the comparisons showed statistically significant differences in spiking during control vs. ES delivery, as assessed through a two-sided paired t-test (with Bonferroni correction for multiple comparisons, not significant: $p > 0.05$). Light colored bars: control; dark colored bars: during ES delivery. Pyramidal: N=21 cells for 8, 30 Hz, N=13 for 140 Hz; Pvalb: N=22 cells for 8, 30 Hz, N=12 for 140 Hz; SST: N=13 for 8, 30 Hz, N=10 for 140 Hz. Bars indicate mean, error bars indicate st.d.

Figure S2

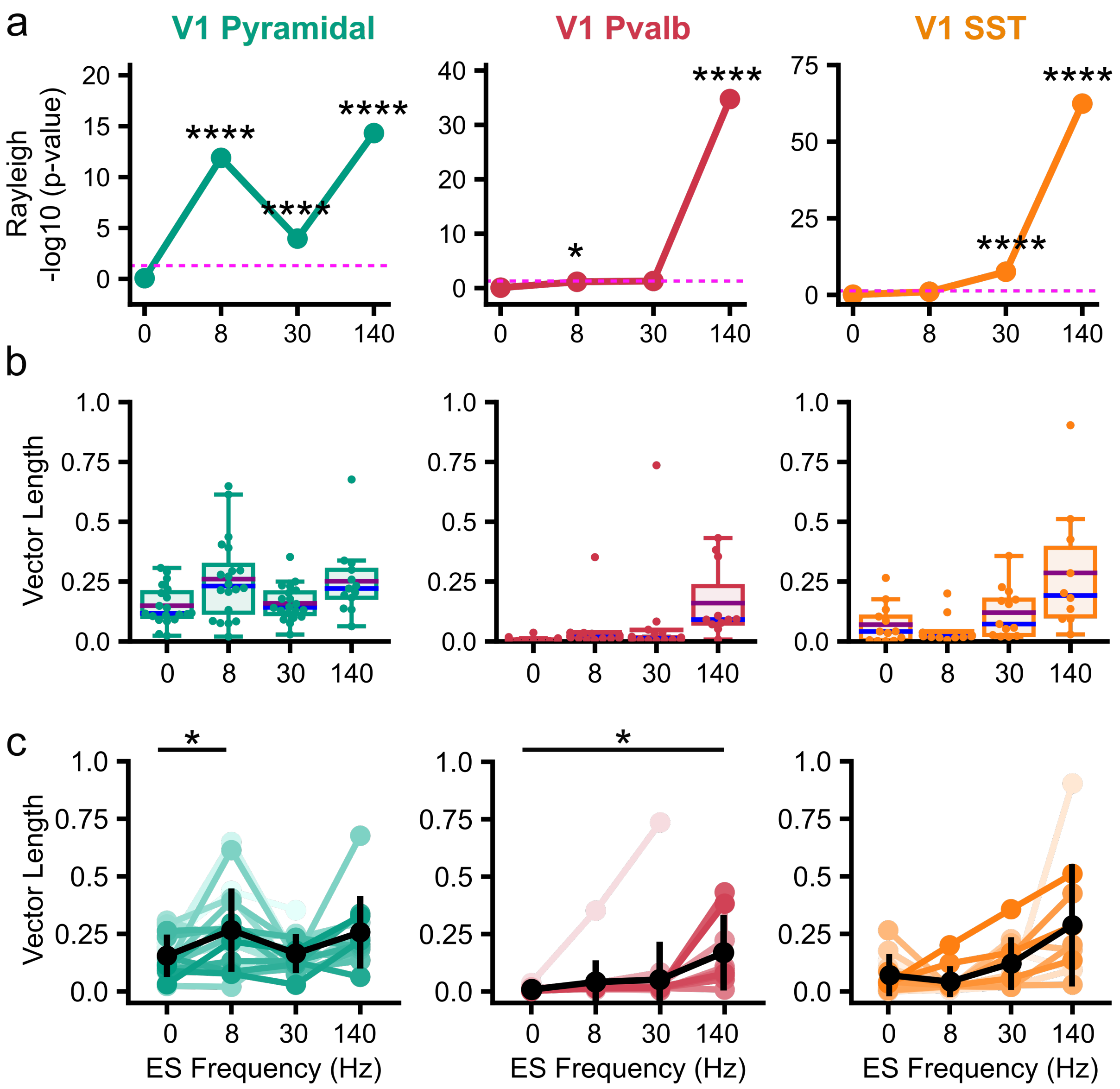


**Figure S2. Summary statistics for ES stimulation at 100 nA in V1 cortex.**
**a-c,** Summary statistics for the second highest ES amplitude tested (100 nA) in V1 classes, against varying ES frequencies (0 (Control, no ES), 8, 30, 140 Hz). **a**, Spike entrainment to ES assessed via the Rayleigh's test (not significant: $p > 0.05$, *: $p < 0.05$, ****: $p < 0.0001$) Dashed pink line in **a**, p=0.05. **b,** The population vector length plotted for every cell in each class (green circles for Pyramidal, red for Pvalb, orange for SST). Boxplots shows the quartiles of the dataset, whiskers: rest of the distribution. Purple and blue lines: mean and median, respectively. **c,** The population vector length for each cell (from **b**) across ES frequencies (colored lines). Circles: mean; Error bars: st.d (black). The population vector length is compared against control to assess degree of entrainment (paired t-test, false discovery rate (FDR)-corrected for multiple comparisons: *: p<0.05, **: p<0.01, see Table S2). Pyramidal: N=21 cells for 8, 30 Hz, N=13 for 140 Hz; Pvalb: N=22 cells for 8, 30 Hz, N=12 for 140 Hz; SST: N=13 for 8, 30 Hz, N=10 for 140 Hz.

**Figure S3**

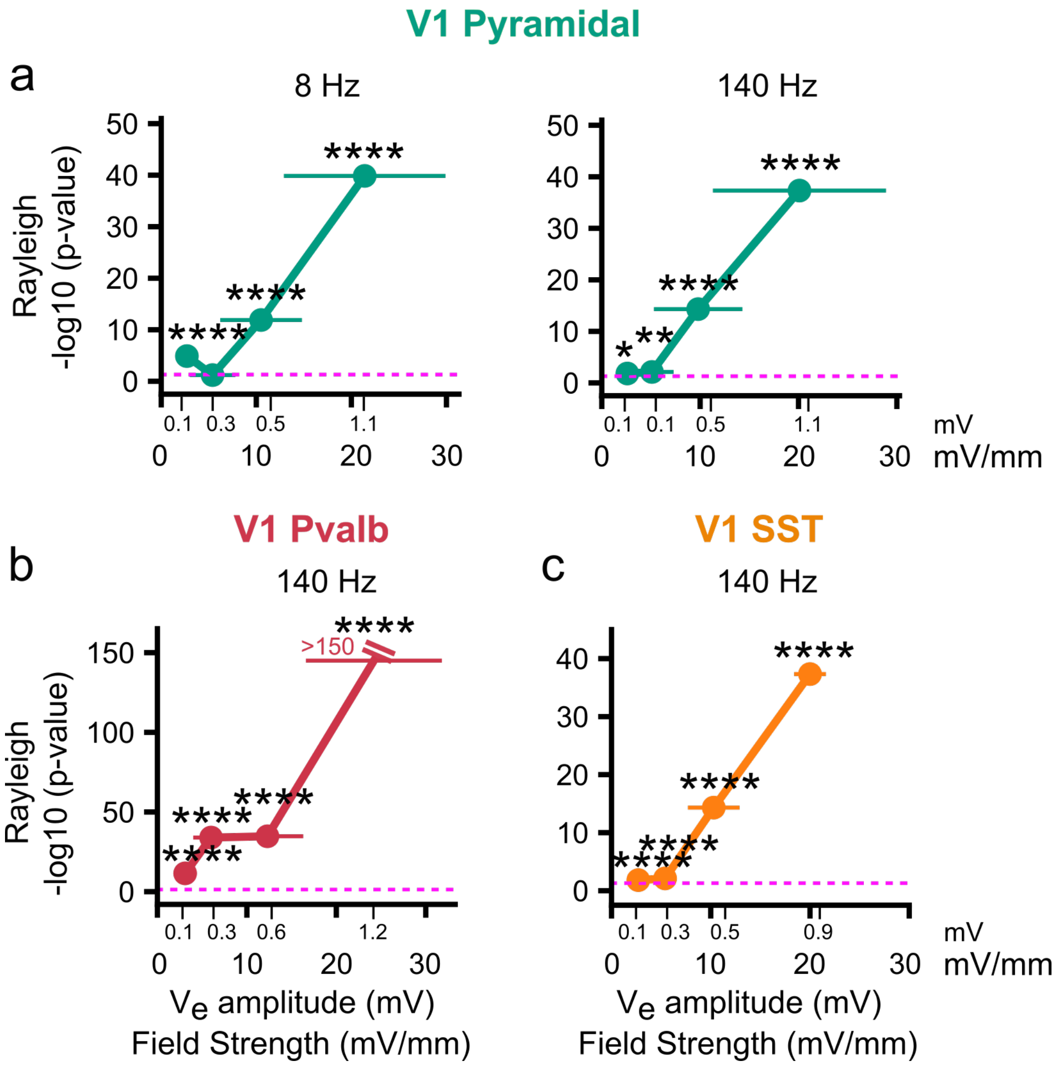


**Figure S3. Strong and robust spike-field entrainment occurs already at the lowest ES amplitude across cell classes. a-c,** p-values calculated for pyramidal, Pvalb, SST cell spikes (left to right) via Rayleigh's test as function of field strength generated by the applied ES amplitudes (at 25, 50, 100, 200 nA, as in Fig 3c). Plotted here are the ES frequencies showing the strongest entrainment in Figure 3c. All cell classes show statistically significant spike phase entrainment (*p < 0.05, **:p < 0.01, ***p < 0.001, ****p < 0.0001) beginning even at the lowest ES amplitudes applied our study. Dashed pink line indicates p-value at 0.05. Pyramidal: N=21 cells for 8 Hz, N=13 for 140 Hz; Pvalb: N=12 for 140 Hz; SST: N=10 for 140 Hz.

**Figure S4**

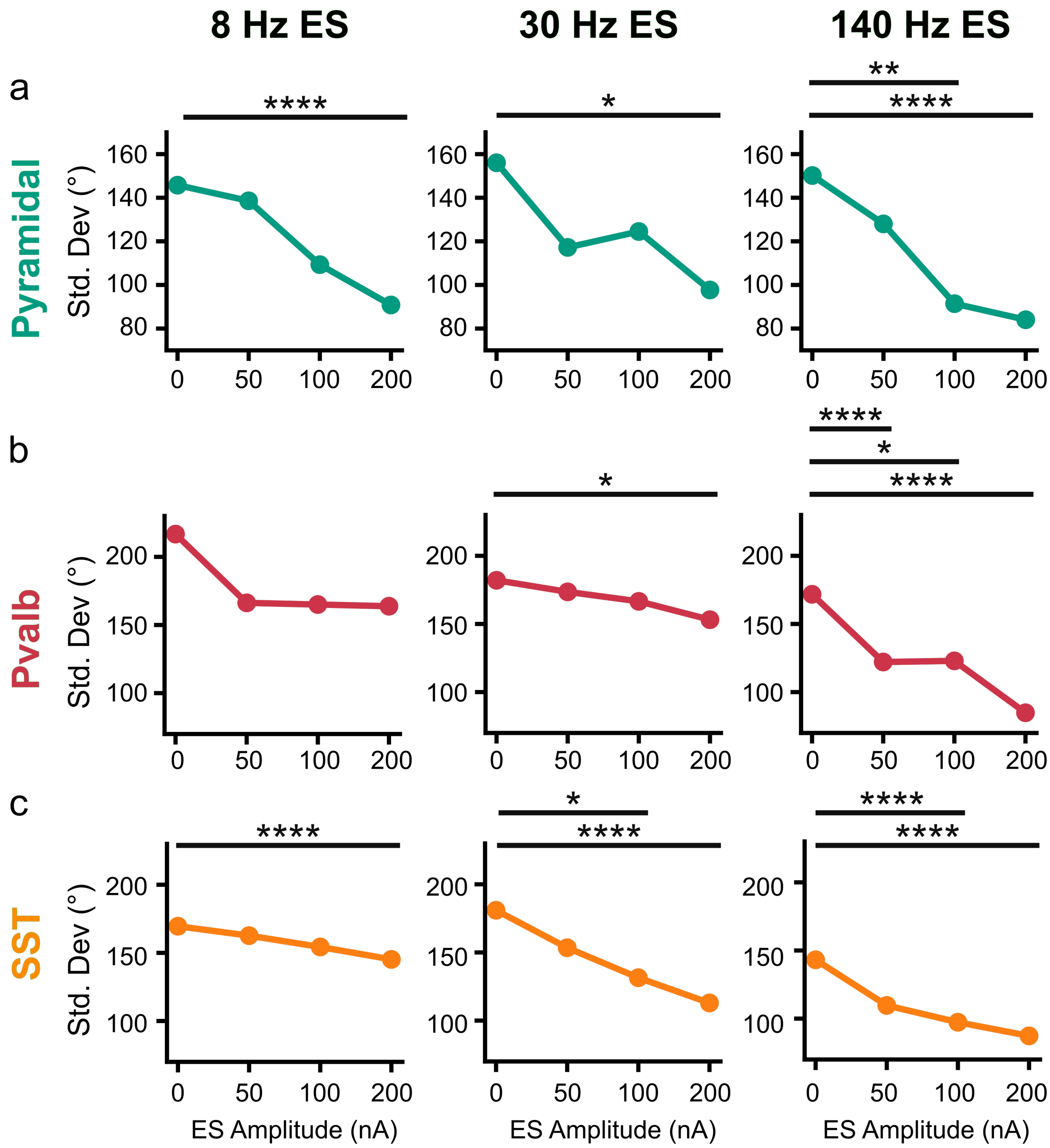


**Figure S4. Spike-phase distribution variability decreases as ES Amplitude is increased. a-c,** Circular standard deviation of the spike-phase distribution across ES frequencies (8, 30, 140 Hz) at increasing ES amplitudes (0, 50, 100, 200 nA), for Pyramidal (green), Pvalb (red), and SST (orange) classes. The standard deviation after ES is compared against control to assess the variabiltiy in the spike-phase distribution. (Levene's test, false discovery rate (FDR)-corrected for multiple comparisons: not significant: $p > 0.05$, *: $p < 0.05$, **: $p<0.01$, ****: $p<0.0001$, see Table S2). Pyramidal: N=21 cells for 8, 30 Hz, N=13 for 140 Hz; Pvalb: N=22 cells for 8, 30 Hz, N=12 for 140 Hz; Sst: N=13 for 8, 30 Hz, N=10 for 140 Hz.

**Figure S5**

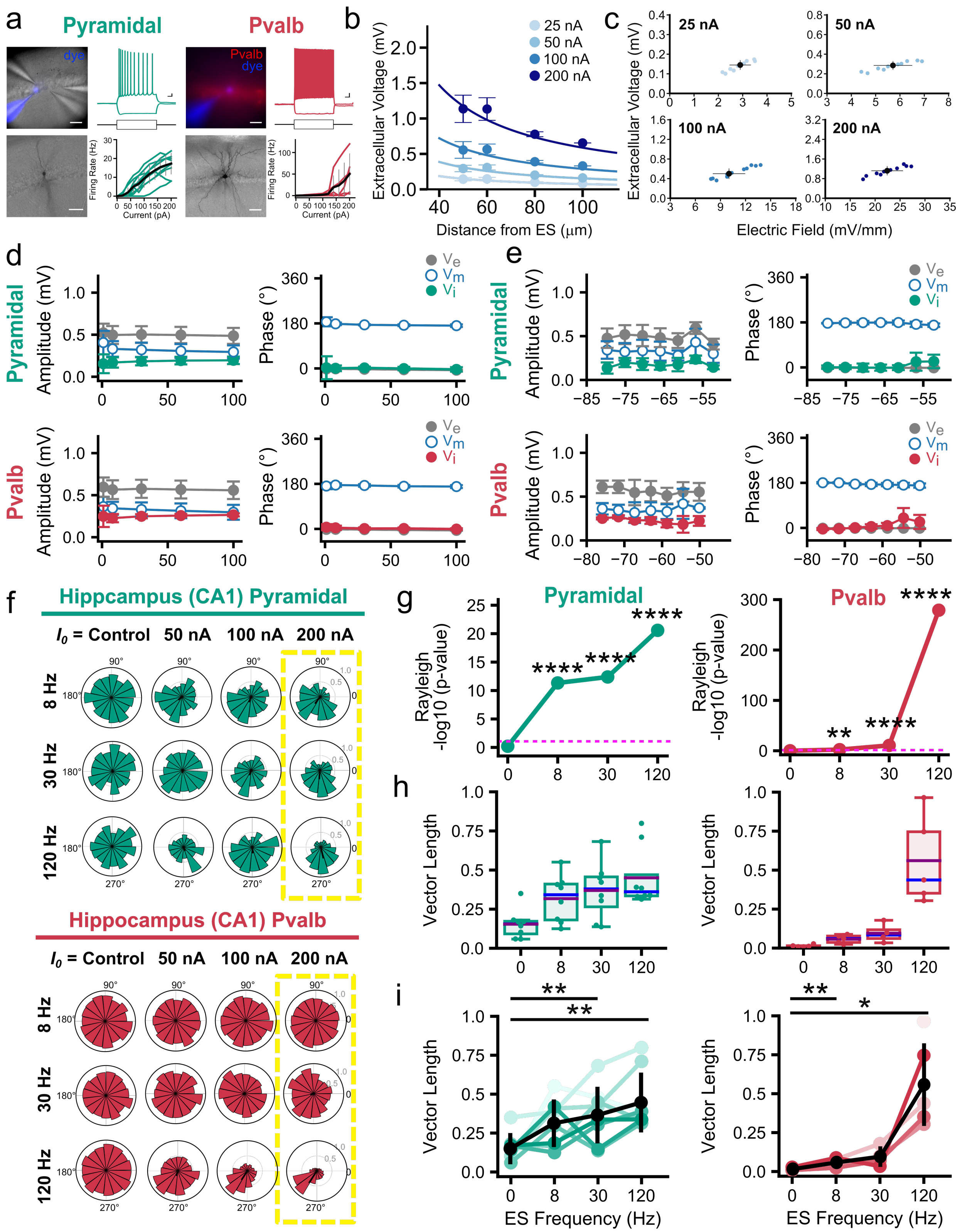
a
Pyramidal
Pvalb
dye
Pvalb
dye
Firing Rate (Hz)
Current (pA)
b
Extracellular Voltage (mV)
Distance from ES (μm)
25 nA
50 nA
100 nA
200 nA
c
Extracellular Voltage (mV)
25 nA
50 nA
100 nA
200 nA
Electric Field (mV/mm)
d
Pyramidal
Pvalb
Amplitude (mV)
Phase (°)
$V_e$
$V_m$
$V_i$
e
Pyramidal
Pvalb
Amplitude (mV)
Phase (°)
f
Hippcampus (CA1) Pyramidal
$I_0$ = Control
50 nA
100 nA
200 nA
8 Hz
30 Hz
120 Hz
Hippocampus (CA1) Pvalb
g
Pyramidal
Pvalb
Rayleigh
-log10 (p-value)
****
**
h
Vector Length
i
Vector Length
ES Frequency (Hz)
**
*

**Figure S5. ES entrainment of sub- and supra-threshold activity is present in subcortical hippocampal regions. a**, (left, Pyramidal; right, Pvalb) Dye-filled recorded cells identified per the CA1 pyramidal layer (pyramidal) or fluorescent marker (Pvalb), cellular morphology (revealed via biocytin imaging), and electrophysiology responses to hyperpolarzing (-140 pA) and depolarizing (100pA for Pyramidal; 300 pA for Pvalb) current injections, and resulting f-I curves from hippocampal pyramidal and Pvalb cells. **b,** Left: $V_e$ amplitude as function of distance between ES and recording electrodes (ES: sinusoidal stimulus amplitude is 25-200 pA; ES frequency: 8 Hz; circles: $V_e$ mean amplitude; error bars: $V_e$ amplitude st.d.) Trendlines are least-squares fit of the point source approximation. **c**, $V_e$ and electric field amplitude induced by the ES at the extracellular recording electrode closest to the whole-cell patched soma (approx. 15 µm). Blue: $V_e$ amplitude for each experiment (n=10); Black: mean and st.d. across experiments. **d, e,** Top: CA1 pyramidal neurons, Bottom: CA1 Pvalb. ES effect on neurons at resting potential (d) and at hyper- and depolarized potentials (e), where neurons were held at a range of membrane potentials via injection of depolarizing or hyperpolarizing current $I_{inj}$ (from -90 to 90 pA). (ES amplitude: 100 nA, frequency: 1 to 100 Hz). ES Amplitude (top) and phase (bottom) with $V_e$ (gray), $V_i$ (green for Pyramidal, red for Pvalb), and $V_m$ (blue outlined circle). Circles: mean; error bars: st.d. Hippocampal neurons also exhibit ES-frequency independence with induced $V_i$, $V_e$ and $V_m$ amplitude and phase remaining constant for ES frequencies ranging 1-100 Hz. The induced ES effects on $V_i$, $V_e$ and $V_m$ amplitude and phase are also broadly independent of membrane polarization. N=9 cells for Pyramidal, N=6 for Pvalb cell classes. **f,** Spike-phase distribution for the hippocampal pyramidal (top, green) and Pvalb (bottom, red) neurons. (Rows) ES frequency (top to bottom): 8, 30, to 120 Hz. (Columns) ES amplitude (left to right): 0 (Control), 50, 100 and 200 nA. Control: no ES (ES amplitude: 0 nA). Increasing (but subthreshold) Pyramidal cells entrain across ES frequencies whereas Pvalb exhibit selective entrainment to high ES frequencies. **g-i,** Summary statistics for the highest ES amplitude (200 nA, yellow box in f) against ES frequencies: 0 (control, no ES), 8, 30, 120 Hz. **g**, Spike entrainment to ES assessed via the Rayleigh's test (not significant: $p > 0.05$, *: $p < 0.05$, **: $p < 0.01$, ***: $p < 0.001$, see Table S3). Dashed pink line in **g**: p=0.05. **h,** The population vector length plotted for each cell across cell classes (green circles: Pyramidal; red: Pvalb). Boxplots shows the quartiles of the dataset, while the whiskers show the rest of the distribution (purple line: mean; blue lines: median). **i,** The population vector length for each cell (from **h**) across ES frequencies (colored lines) and mean value with st.d. (black line). The population vector length during each ES frequency is compared against control to assess degree of entrainment (paired t-test, false discovery rate (FDR)-corrected for multiple comparisons: *: $p<0.05$, **: $p<0.01$, see Table S3). Pyramidal: N=8 cells, Pvalb: N=5 cells.

**Figure S6**

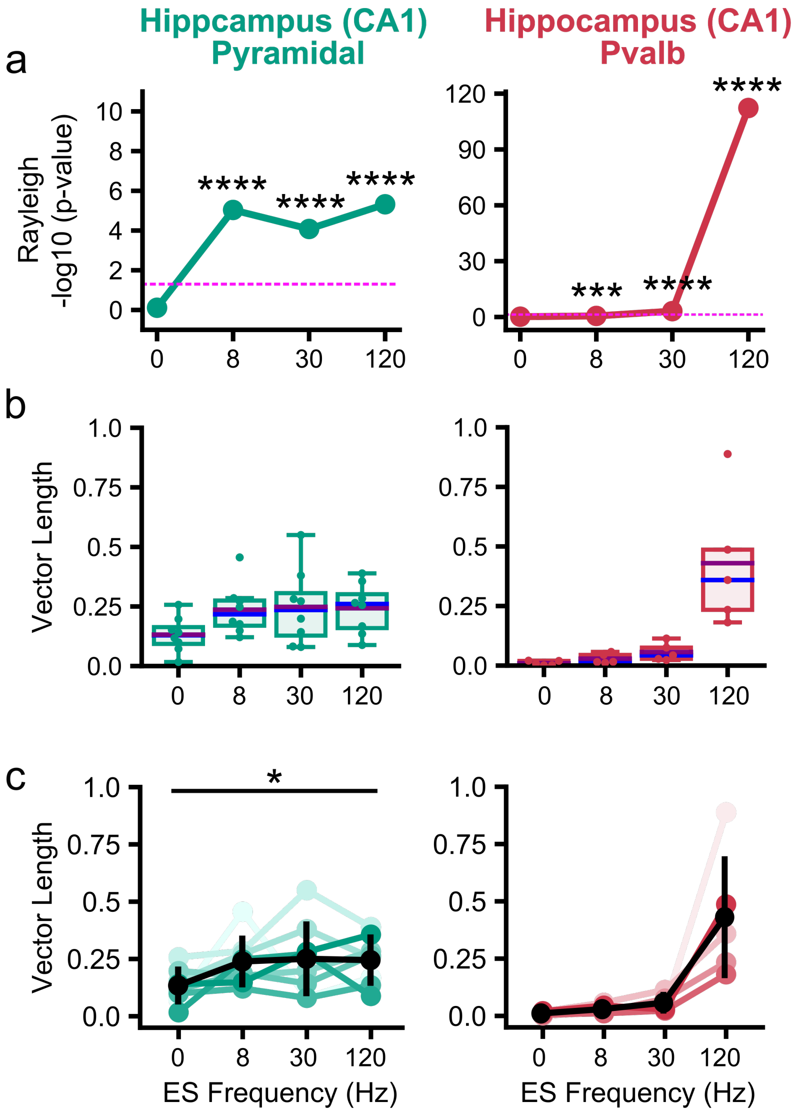


**Figure S6. Summary statistics for ES stimulation at 100 nA in CA1 hippocampus. a-c,** Summary statistics for the second highest ES amplitude tested (100 nA) in CA1 against varying ES frequencies: 0 (control, no ES), 8, 30, 120 Hz. Spike entrainment assessed via the Rayleigh's test (not significant, n.s.: $p > 0.05$, *$p < 0.05$, **:$p < 0.01$, ***$p < 0.001$). Dashed pink line in **a**: $p$=0.05. **b,** The population vector length plotted for each cell in each class (green circles for Pyramidal, red for Pvalb). Boxplots shows the quartiles of the dataset while the whiskers extend to show the rest of the distribution. **c,** The population vector length across ES frequencies (colored lines) and mean value with st.d. (black line). The population vector length during each ES frequency is compared against control to assess degree of entrainment (paired t-test, false discovery rate (FDR)-corrected for multiple comparisons: *: $p<0.05$, see Table S3). Pyramidal: N=8 cells, Pvalb: N=5 cells.

**Figure S7**

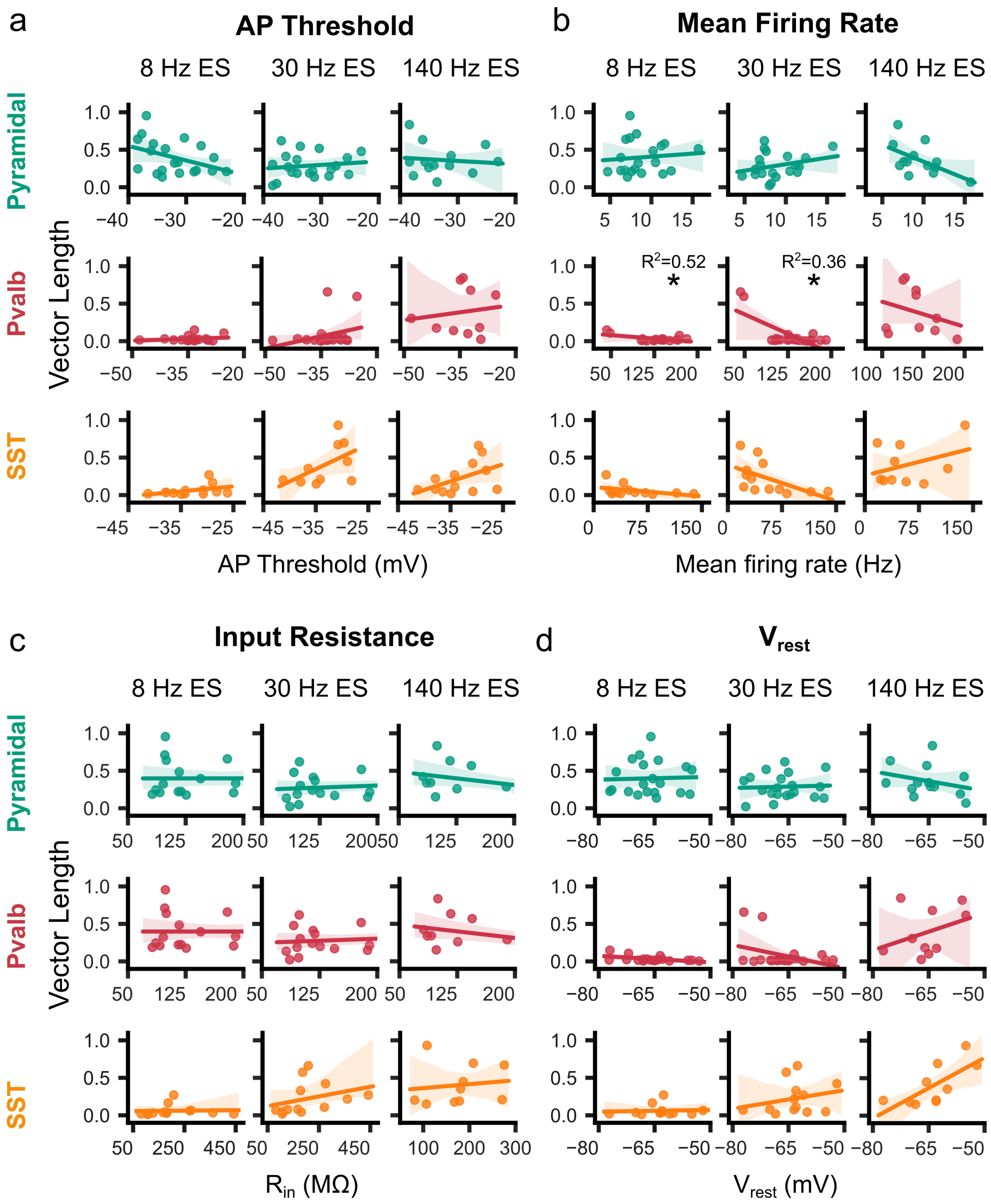
a
AP Threshold
b
Mean Firing Rate
8 Hz ES
30 Hz ES
140 Hz ES
Pyramidal
Pvalb
SST
Vector Length
R²=0.52
R²=0.36
*
AP Threshold (mV)
Mean firing rate (Hz)
c
Input Resistance
d
V_rest
Rin (MΩ)
Vrest (mV)

**Figure S7. Population vector length does not correlate with cellular intrinsic properties.** Several cellular intrinsic properties examined for correlations with the degree of ES entrainment (assessed via population vector length). The vector length as a function of **a,** AP (action potential) threshold, **b,** mean firing rate, **c,** input Resistance, and **d,** $V_{rest}$ (resting membrane potential) during ES applied at various frequencies (8, 30, 140 Hz ES). Circles: cells in Pyramidal (green), Pvalb (red), and SST (orange) cell classes (ES amplitude: 200 nA). Linear regression (line) and 95% confidence intervals (shaded area). (Not significant, ns): $p > 0.01$,*: $p<0.01$). Pyramidal: N=21 cells for 8, 30 Hz, N=13 for 140 Hz; Pvalb: N=22 cells for 8, 30 Hz, N=12 for 140 Hz; SST: N=13 for 8, 30 Hz, N=10 for 140 Hz.

### Figure S8

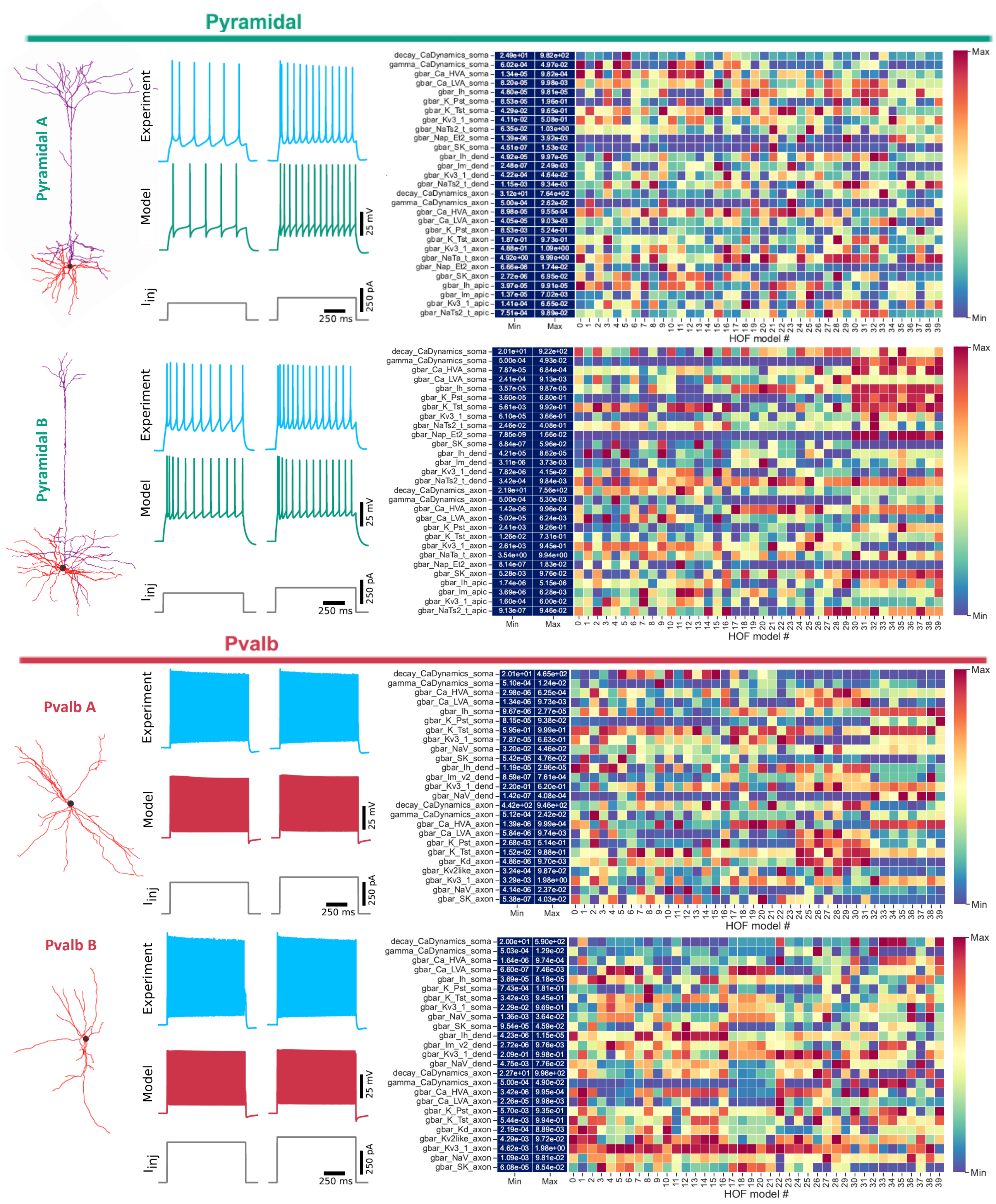

Pyramidal
Pyramidal A
Pyramidal B
Experiment
Model
Iinj
250 ms
25 mV
250 pA
Min
Max
HOF model #
decay_CaDynamics_soma
gamma_CaDynamics_soma
gbar_Ca_HVA_soma
gbar_Ca_LVA_soma
gbar_Ih_soma
gbar_K_Pst_soma
gbar_K_Tst_soma
gbar_Kv3_1_soma
gbar_NaTs2_t_soma
gbar_Nap_Et2_soma
gbar_SK_soma
gbar_Ih_dend
gbar_Im_dend
gbar_Kv3_1_dend
gbar_NaTs2_t_dend
decay_CaDynamics_axon
gamma_CaDynamics_axon
gbar_Ca_HVA_axon
gbar_Ca_LVA_axon
gbar_K_Pst_axon
gbar_K_Tst_axon
gbar_Kv3_1_axon
gbar_NaTa_t_axon
gbar_Nap_Et2_axon
gbar_SK_axon
gbar_Ih_apic
gbar_Im_apic
gbar_Kv3_1_apic
gbar_NaTs2_t_apic
Pvalb
Pvalb A
Pvalb B
gbar_NaV_soma
gbar_Im_v2_dend
gbar_NaV_dend
gbar_Kd_axon
gbar_Kv2like_axon
gbar_NaV_axon

**Figure S8. Multi-objective evolutionary optimization produces cell class-specific excitatory and inhibitory single-cell models.** Left, reconstructed morphology of single-cell neurons (top: two layer 5 pyramidal neurons; bottom: two layer 5 Pvalb neurons). Middle, experimental (experiment) and simulation (model) intracellular voltage traces for two different injected DC current stimuli ($I_{inj}$). Right, conductance parameter distribution across the 40 hof models for each cell (y-axis: individual conductances; blue columns: min-max values for each conductance across hof models; colors: value for each conductance for each hof model; x-axis: hof model # of each cell) The 40 best hof models per cell are considered (hof model # 0-39). The multi-objective evolutionary optimization results in a hof models with a distribution of conductance values. Hof model #0 is the model with the best performance across the population of 40 for matching a set electrophysiology features on intracellular current stimuli they trained on (dc inputs; training set) [40]. In general, hof models generalize equally well for stimuli they did not train on (noise inputs; validation set) [40].

**Figure S9**

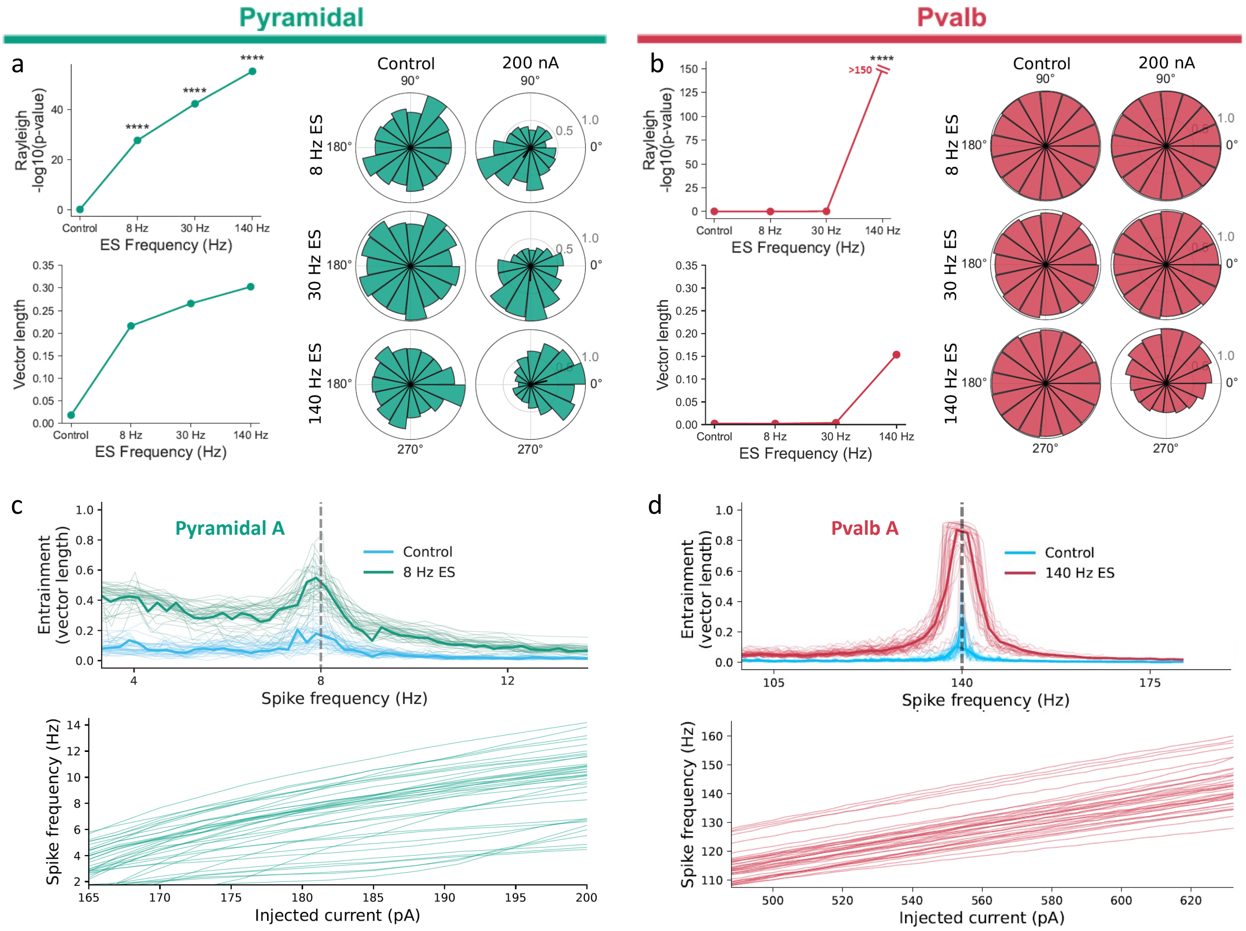

Pyramidal
Pvalb
a
b
c
d
Rayleigh -log10(p-value)
ES Frequency (Hz)
Vector length
Control
200 nA
8 Hz ES
30 Hz ES
140 Hz ES
>150
Pyramidal A
Pvalb A
Control
8 Hz ES
140 Hz ES
Entrainment (vector length)
Spike frequency (Hz)
Injected current (pA)

**Figure S9. Computational investigation of mechanisms affecting the entrainment of spiking to ES. a, Top left,** p-values calculated via Rayleigh's test (null: spike sample drawn from a homogeneous distribution). **a, Bottom left;** spike-phase coupling as measured via the population vector length. **a, Right side,** Spike-phase distributions for different ES parameters vs. control. (Rows) ES frequency (top to bottom): 8, 30, to 140 Hz. (Columns) ES amplitude (left to right): 0 (Control) and 200 nA. "Control" indicates no ES (ES amplitude: 0 nA). Green: Pyramidal cell A, hof model # 0. **b**, Same as panel for Pvalb model A, hof 0. **c, Top,** Spike-phase coupling strength (population vector length) for ES 8 Hz and 200 nA as function of spike frequency, to assess the relationship between spike frequency and entrainment as in Fig. 4. Each line represents one of 40 different hof models for the same morphology. Thick line: the median across hof models. "Control" indicates ES amplitude of 0 nA. **c, Bottom,** Spike rate response of each of the hof models of Pyramidal model A for varying intracellular DC current injection amplitude. d, same as panel c but for hof models of Pvalb model A. **d**, Same layout as panel c but for the Pvalb model A across hof models.

**Figure S10**

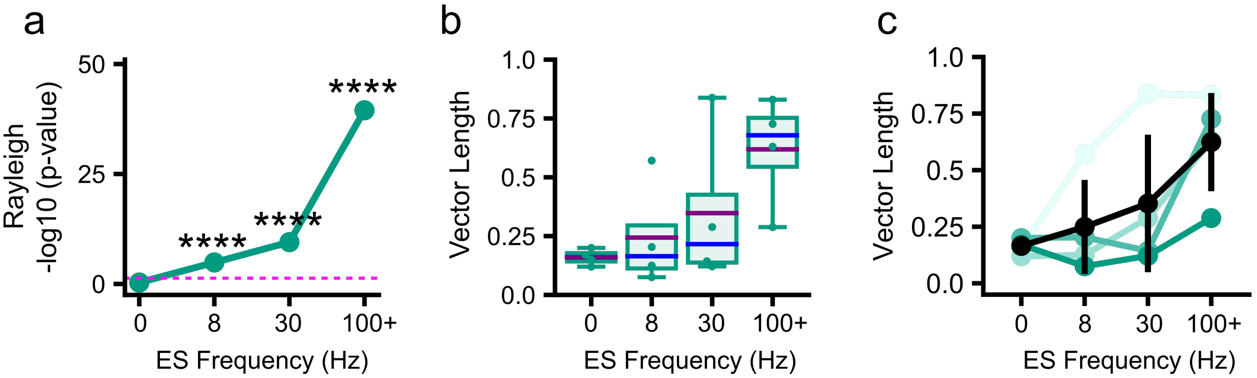


**Figure S10. Summary statistics for ES stimulation at 100 nA in human cortex.**
**a-c,** Summary statistics for the second highest ES amplitude tested in human neurons (100 nA) against varying ES frequencies (0 (Control, no ES), 8, 30, 100+ Hz). **g**, Spike entrainment to ES assessed via the Rayleigh's test. (****p < 0.001) Dashed pink line in **a** indicates p-value at 0.05. b**,** The population vector length plotted for each cell the human Pyramidal cell class (green circles). Boxplots shows the quartiles of the dataset while the whiskers extend to show the rest of the distribution. Purple and blue lines indicate mean and median respectively. c**,** The population vector length for each cell (from **c**) across ES frequencies (colored lines) and mean value with st.d (black line). The population vector length during each ES frequency is compared against Control conditions to assess degree of entrainment. entrainment (paired t-test, false discovery rate (FDR)-corrected for multiple comparisons, not significant: p > 0.05). N=4 human pyramidal cells.

**Table S1**

| | **Pyramidal** | **Pvalb** | **SST** |
|---|---|---|---|
| Rheobase (pA) | 70.0 ± 33.4 | 211.4 ± 76.4 | 88.5 ± 87.0 |
| Resting membrane potential (mV) | -65.5 ± 6.3 | -66.5 ± 6.2 | -63.5 ± 6.3 |
| AP threshold (mV) | -32.2 ± 4.6 | -33.7 ± 4.8 | -33.2 ± 4.5 |
| Induced firing rate (Hz) | 8.5 ± 3.0 | 133.38 ± 35.7 | 57.4 ± 37.8 |
| Input resistance (MΩ) | 157.0 ± 74.6 | 123.2 ± 27.5 | 207.4 ± 105.4 |

**Summary table of mean cellular properties of recorded neurons in V1 cell classes.** Properties were calculated after intracellular current injection, as described in Methods. Data are presented as mean ± std. Pyramidal: N=24 cells, Pvalb: N=22 cells, Sst: N=13.

**Table S2**

| Pyramidal | | Control | 50 nA | 100 nA | 200 nA |
|---|---|---|---|---|---|
| 8 Hz | # of spikes<br>Rayleigh p-value<br>vector length<br>vector angle<br>angle st.d.<br>kappa | n=1033<br>p=0.50<br>0.03<br>285.46<br>145.75<br>0.07 | n=959<br>p=6.3e-02<br>0.05<br>251.10<br>138.58<br>0.11 | n=1036<br>p=1.3e-12<br>0.16<br>258.79<br>109.29<br>0.33 | n=1108<br>p=1.4e-40<br>0.29<br>256.31<br>90.80<br>0.59 |
| 30 Hz | # of spikes<br>Rayleigh p-value<br>vector length<br>vector angle<br>angle st.d.<br>kappa | n=1003<br>p=0.85<br>0.01<br>204.11<br>156.04<br>0.03 | n=986<br>p=2.9e-07<br>0.12<br>299.83<br>117.22<br>0.24 | n=1028<br>p=1.1e-04<br>0.09<br>259.33<br>124.50<br>0.19 | n=994<br>p=1.3e-24<br>0.23<br>262.13<br>97.69<br>0.49 |
| 140 Hz | # of spikes<br>Rayleigh p-value<br>vector length<br>vector angle<br>angle st.d.<br>kappa | n=727<br>p=0.56<br>0.02<br>238.09<br>150.18<br>0.04 | n=716<br>p=7.6e-03<br>0.08<br>274.60<br>127.99<br>0.16 | n=748<br>p=4.8e-15<br>0.21<br>255.09<br>101.40<br>0.42 | n=717<br>p=4.7e-38<br>0.34<br>249.23<br>84.03<br>0.72 |
| **Pvalb** | | **Control** | **50 nA** | **100 nA** | **200 nA** |
| 8 Hz | # of spikes<br>Rayleigh p-value<br>vector length<br>vector angle<br>angle st.d.<br>kappa | n=11536<br>p=0.07<br>0.01<br>266.87<br>216.69<br>0.01 | n=11836<br>p=0.07<br>0.01<br>286.04<br>166.24<br>0.03 | n=10678<br>p=0.07<br>0.02<br>272.06<br>165.02<br>0.03 | n=11100<br>p=0.043<br>0.02<br>263.51<br>163.77<br>0.03 |
| 30 Hz | # of spikes<br>Rayleigh p-value<br>vector length<br>vector angle<br>angle st.d.<br>kappa | n= 13920<br>p=0.85<br>0.01<br>265.15<br>182.04<br>0.002 | n=14373<br>p=0.23<br>0.01<br>262.82<br>173.5400021<br>0.02 | n=14115<br>p=0.05<br>0.01<br>234.12<br>166.5671075<br>0.03 | n=13572<br>p=2.2e-05<br>0.03<br>228.02<br>153.116115<br>0.06 |
| 140 Hz | # of spikes<br>Rayleigh p-value<br>vector length<br>vector angle<br>angle st.d.<br>kappa | n=7273<br>p=0.29<br>0.01<br>288.14<br>171.78<br>0.01 | n=7305<br>p=1.1e-34<br>0.1<br>225.12<br>122.08<br>0.20 | n=7951<br>p=1.8e-35<br>0.1<br>214.41<br>122.91<br>0.20 | n=7502<br>p=0<br>0.33<br>216.78<br>84.83<br>0.71 |
| **SST** | | **Control** | **50 nA** | **100 nA** | **200 nA** |
| 8 Hz | # of spikes<br>Rayleigh p-value<br>vector length<br>vector angle<br>angle st.d. | n=3459<br>p=0.54<br>0.01<br>265.35<br>169.53 | n=3580<br>p=3.2e-01<br>0.02<br>263.67<br>162.64 | n=3347<br>p=9.4e-02<br>0.03<br>237.07<br>154.32 | n=3451<br>p=3.7e-03<br>0.04<br>221.33<br>145.20 |

| | | | | | |
|---|---|---|---|---|---|
| | kappa | 0.007 | 0.03 | 0.05 | 0.08 |
| 30 Hz | # of spikes | n=3455 | n=3608 | n=3357 | n=3400 |
| | Rayleigh p-value | p=0.861 | p=6.3e-02 | p=2.8e-08 | p=3.0e-31 |
| | vector length | 0.01 | 0.03 | 0.07 | 0.14 |
| | vector angle | 208.68 | 255.53 | 233.78 | 221.42 |
| | angle st.d. | 180.95 | 153.43 | 131.45 | 112.92 |
| | kappa | 0.00 | 0.06 | 0.14 | 0.29 |
| 140 Hz | # of spikes | n=1311 | n=2412 | n=2538 | n=3045 |
| | Rayleigh p-value | p=0.330 | p=9.3e-28 | p=3.9e-63 | p=1.4e-133 |
| | vector length | 0.02 | 0.16 | 0.24 | 0.31 |
| | vector angle | 198.09 | 220.90 | 231,51 | 224.28 |
| | angle st.d. | 143.16 | 109.66 | 97.33 | 87.33 |
| | kappa | 0.02 | 0.32 | 0.48 | 0.66 |

**Output values of circular statistics tests conducted for suprathreshold entrainment analysis of V1 cell classes in Figure 3b.** Pyramidal (Pyr): N=21 cells for 8, 30 Hz, N=13 for 140 Hz; Pvalb: N=22 cells for 8, 30 Hz, N=12 for 140 Hz; SST: N=13 for 8, 30 Hz, N=10 for 140 Hz. (st.d.: standard deviation)

**Table S3**

| Pyramidal | | Control | 50 nA | 100 nA | 200 nA |
|---|---|---|---|---|---|
| 8 Hz | # spikes<br>Rayleigh p-value<br>vector length<br>vector angle<br>kappa | n=336<br>p=0.84<br>0.02<br>291.31<br>0.05 | n=326<br>p=5.8e-04<br>0.15<br>219.52<br>0.31 | n=344<br>p=9.2e-06<br>0.18<br>248.05<br>0.37 | n=339<br>p=4.6e-12<br>0.28<br>231.71<br>0.57 |
| 30 Hz | # spikes<br>Rayleigh p-value<br>vector length<br>vector angle<br>kappa | n=328<br>p=0.77<br>0.03<br>356.6<br>0.06 | 333<br>p=0.11<br>0.08<br>268.73<br>0.16 | 326<br>p=8.34-05<br>0.17<br>246.98<br>0.34 | 324<br>p=4.1e-13<br>0.29<br>265.85<br>0.61 |
| 140 Hz | # spikes<br>Rayleigh p-value<br>vector length<br>vector angle<br>kappa | n=320<br>p=0.62<br>0.04<br>312.76<br>0.08 | 331<br>p=3.3e-03<br>0.13<br>290.39<br>0.26 | 306<br>p=4.8e-06<br>0.2<br>275.73<br>0.41 | 323<br>p=2.6e-21<br>0.38<br>282.77<br>0.81 |
| **Pvalb** | | **Control** | **50 nA** | **100 nA** | **200 nA** |
| 8 Hz | # spikes<br>Rayleigh p-value<br>vector length<br>vector angle<br>kappa | 2045<br>0.802<br>0.01<br>270<br>0.02 | 2092<br>0.106<br>0.03<br>306.93<br>0.07 | 1832<br>0.27<br>0.03<br>260.52<br>0.05 | 2212<br>p=5.0e-03<br>0.05<br>241.5<br>0.1 |
| 30 Hz | # spikes<br>Vector Length<br>p_values<br>Vector Angle<br>kappa | 1929<br>p=0.692<br>0.01<br>238.06<br>0.03 | 1958<br>0.329<br>0.02<br>271.91<br>0.05 | 1779<br>p=7.2e-04<br>0.06<br>199.42<br>0.13 | 2050<br>p=2.3e-11<br>0.11<br>197.28<br>0.22 |
| 140 Hz | # spikes<br>Vector Length<br>p_values<br>Vector Angle<br>kappa | 2274<br>p=0.528<br>0.02<br>318.14<br>0.03 | 2050<br>p=5.5e-25<br>0.16<br>254.56<br>0.33 | 2052<br>p=5.3e-113<br>0.35<br>237.14<br>0.75 | 2721<br>p=1.3e-279<br>0.47<br>236.89<br>1.07 |

**Output values of circular statistics tests conducted for suprathreshold entrainment analysis of hippocampal CA1 cell classes in Figure S5.** Pyramidal: N=8 cells, Pvalb: N=5 cells.

**Table S4**

| Pyramidal | | Control | 50 nA | 100 nA | 200 nA |
|---|---|---|---|---|---|
| 8 Hz | # spikes | n=236 | n=233 | n=231 | n=245 |
| | Rayleigh p-value | p=0.49 | p=3.7e-05 | p=1.3e-05 | p=7.9e-20 |
| | vector length | 0.02 | 0.15 | 0.18 | 0.28 |
| | vector angle | 291.31 | 219.52 | 248.05 | 231.71 |
| | kappa | 0.05 | 0.31 | 0.37 | 0.57 |
| 30 Hz | # spikes | n=236 | n=233 | n=219 | n=257 |
| | Rayleigh p-value | p=0.60 | p=2.3e-07 | P=3.1e-10 | p=2.8e-41 |
| | vector length | 0.05 | 0.25 | 0.31 | 0.58 |
| | vector angle | 273.03 | 252.22 | 244.68 | 231.05 |
| | kappa | 0.09 | 0.55 | 0.66 | 1.41 |
| 140 Hz | # spikes | n=243 | n=238 | n=234 | n=258 |
| | Rayleigh p-value | p=0.6 | p=5.1e-20 | p=3.23e-40 | p=2.3e-64 |
| | vector length | 0.05 | 0.42 | 0.59 | 0.7 |
| | vector angle | 117.32 | 296.26 | 289.39 | 289.31 |
| | kappa | 0.09 | 0.93 | 1.48 | 2 |

**Output values of circular statistics tests conducted for suprathreshold entrainment analysis of human neurons in Figure 6.** Pyramidal: N=4 cells.
